## Supplementary material for "Evidence for deliberate burial of the dead by *Homo naledi*": Table 3 Main Manuscript

| Samples | Li (mg/kg) | Be (mg/kg) | Sc (mg/kg) | Ti (mg/kg) | V (mg/kg) | Cr (mg/kg) | Co (mg/kg) | Ni (mg/kg) |
|---|---|---|---|---|---|---|---|---|
| DF1 | 62.72 | 4.98 | 13.72 | 2857.59 | 29.82 | 71.33 | 17.34 | 78.89 |
| DF3 | 97.31 | 3.64 | 24.69 | 6752.72 | 125.06 | 190.01 | 13.90 | 59.81 |
| DF5 | 66.58 | 4.55 | 13.21 | 2671.05 | 27.56 | 58.30 | 19.16 | 65.78 |
| DF7 | 93.91 | 4.63 | 21.61 | 4339.39 | 50.68 | 106.04 | 22.37 | 86.51 |
| DF9 | 54.05 | 1.40 | 9.39 | 2707.41 | 55.68 | 88.01 | 19.60 | 65.77 |
| DF11 | 71.00 | 2.05 | 12.27 | 3302.62 | 65.67 | 104.88 | 20.76 | 66.81 |
| DF13 | 61.63 | 2.43 | 11.86 | 2891.16 | 67.41 | 102.01 | 19.74 | 69.90 |
| DF15 | 72.00 | 4.84 | 17.94 | 3892.83 | 59.89 | 111.77 | 21.85 | 79.72 |
| SA1 | 195.19 | 2.82 | 22.27 | 5582.73 | 116.10 | 179.88 | 31.24 | 86.39 |
| SA2 | 54.83 | 3.03 | 11.81 | 2180.48 | 25.29 | 63.55 | 12.30 | 62.15 |
| SA3 | 151.79 | 4.03 | 17.30 | 4936.23 | 74.00 | 138.72 | 26.25 | 89.05 |
| SA4 | 31.73 | 3.22 | 7.32 | 879.79 | 11.99 | 27.58 | 8.76 | 46.68 |
| SA5 | 41.46 | 3.96 | 9.12 | 1126.62 | 10.97 | 35.99 | 11.91 | 61.94 |
| SA6 | 121.89 | 4.91 | 17.94 | 4172.23 | 43.44 | 109.37 | 25.04 | 96.35 |
| SA7 | 163.02 | 4.95 | 21.18 | 5605.44 | 65.66 | 141.74 | 30.99 | 104.83 |
| SA8 | 101.94 | 5.30 | 22.77 | 4559.94 | 50.97 | 122.46 | 21.99 | 90.95 |
| SB1 | 100.00 | 4.54 | 19.74 | 3566.22 | 54.21 | 111.36 | 17.58 | 93.54 |
| SB2 | 99.63 | 4.78 | 20.19 | 3328.53 | 49.91 | 107.87 | 15.41 | 96.18 |
| SB3 | 104.72 | 4.86 | 21.28 | 3735.89 | 48.15 | 112.94 | 18.69 | 96.77 |
| SC1 | 95.33 | 4.87 | 20.13 | 3390.27 | 51.14 | 103.45 | 17.91 | 92.12 |
| SC2 | 99.16 | 5.25 | 21.18 | 3778.99 | 55.08 | 113.74 | 18.85 | 96.56 |
| SC3 | 90.88 | 4.59 | 20.03 | 3415.78 | 49.35 | 105.19 | 17.79 | 85.04 |
| SC4 | 104.65 | 4.77 | 23.16 | 4185.46 | 58.54 | 129.87 | 19.51 | 95.77 |
| SE1 | 109.20 | 1.64 | 9.57 | 2845.81 | 51.67 | 82.85 | 15.17 | 53.81 |
| SE2 | 141.88 | 1.93 | 11.84 | 3374.12 | 51.17 | 99.50 | 17.80 | 65.78 |
| SE3 | 2.24 | 1.11 | 3.10 | 30.67 | 15.40 | 4.92 | 3.64 | 6.83 |
| SE4 | 48.34 | 2.79 | 9.84 | 1973.73 | 28.19 | 58.19 | 12.49 | 56.69 |
| SE5 | 36.18 | 2.67 | 8.63 | 1815.31 | 24.07 | 47.80 | 10.35 | 45.50 |
| FS2280 | 126.71 | 4.96 | 18.55 | 3921.23 | 30.78 | 78.11 | 9.34 | 149.23 |

| Cu (mg/kg) | Zn (mg/kg) | Ga (mg/kg) | Ge (mg/kg) | As (mg/kg) | Se (mg/kg) | Rb (mg/kg) | Sr (mg/kg) | Y (mg/kg) | Zr (mg/kg) |
|---|---|---|---|---|---|---|---|---|---|
| 66.39 | 122.22 | 19.42 | 1.07 | 1.89 | 0.24 | 65.81 | 67.25 | 24.14 | 81.38 |
| 40.73 | 66.53 | 12.92 | 1.48 | 5.51 | 0.25 | 114.51 | 41.20 | 19.54 | 177.90 |
| 63.80 | 135.05 | 15.91 | 1.17 | 1.92 | 0.23 | 63.96 | 60.06 | 32.03 | 75.28 |
| 88.96 | 151.70 | 15.38 | 1.40 | 2.19 | 0.16 | 95.04 | 64.42 | 29.45 | 121.14 |
| 65.51 | 122.42 | 10.37 | 1.05 | 3.30 | 0.16 | 48.53 | 47.39 | 16.63 | 79.72 |
| 84.45 | 137.36 | 10.41 | 1.15 | 3.99 | 0.11 | 64.04 | 36.96 | 21.72 | 96.05 |
| 70.25 | 112.97 | 12.36 | 1.15 | 4.56 | 0.13 | 66.54 | 41.48 | 19.52 | 89.70 |
| 91.66 | 168.75 | 13.75 | 1.28 | 3.96 | 0.19 | 85.49 | 65.16 | 23.05 | 112.46 |
| 70.18 | 70.56 | 12.92 | 1.83 | 7.20 | 0.24 | 100.44 | 28.59 | 22.45 | 156.68 |
| 57.56 | 158.94 | 13.61 | 1.11 | 1.49 | 0.15 | 65.86 | 56.56 | 21.71 | 73.64 |
| 50.83 | 71.61 | 14.04 | 1.53 | 3.48 | 0.19 | 83.04 | 37.40 | 19.73 | 137.22 |
| 48.48 | 93.40 | 12.84 | 0.81 | 1.58 | 0.14 | 46.27 | 82.24 | 22.31 | 42.18 |
| 66.13 | 141.61 | 15.27 | 1.00 | 1.26 | 0.29 | 58.78 | 64.37 | 26.44 | 53.66 |
| 75.18 | 131.03 | 16.53 | 1.30 | 2.80 | 0.23 | 88.74 | 49.28 | 25.85 | 110.11 |
| 64.75 | 106.22 | 19.99 | 1.69 | 2.86 | 0.22 | 119.57 | 49.93 | 28.51 | 143.74 |
| 70.75 | 111.10 | 19.81 | 1.47 | 2.87 | 0.21 | 107.49 | 58.22 | 27.20 | 130.94 |
| 100.64 | 214.49 | 18.87 | 1.49 | 3.63 | 0.34 | 95.99 | 71.09 | 41.22 | 115.86 |
| 113.82 | 277.27 | 19.88 | 1.56 | 2.67 | 0.30 | 81.43 | 65.73 | 39.09 | 117.59 |
| 107.98 | 228.94 | 20.76 | 1.63 | 3.18 | 0.31 | 101.02 | 62.49 | 38.89 | 117.82 |
| 91.29 | 157.33 | 19.46 | 1.57 | 2.51 | 0.32 | 103.27 | 74.26 | 36.58 | 117.86 |
| 92.77 | 173.66 | 19.55 | 1.65 | 3.84 | 0.46 | 105.41 | 68.58 | 36.90 | 122.29 |
| 80.21 | 144.03 | 17.16 | 1.58 | 2.60 | 0.28 | 96.02 | 62.86 | 31.72 | 111.83 |
| 82.25 | 153.67 | 19.75 | 1.54 | 2.51 | 0.32 | 114.85 | 59.57 | 32.74 | 131.98 |
| 40.67 | 46.75 | 6.93 | 1.10 | 2.51 | 0.15 | 48.10 | 36.09 | 12.60 | 76.01 |
| 48.65 | 55.55 | 7.19 | 1.03 | 2.50 | 0.27 | 56.21 | 30.45 | 16.87 | 92.50 |
| 12.69 | 12.97 | 1.91 | 0.21 | 0.94 | 0.13 | 2.14 | 73.80 | 7.03 | 0.95 |
| 46.06 | 75.99 | 12.07 | 0.85 | 1.96 | 0.24 | 44.37 | 44.13 | 17.21 | 55.70 |
| 49.41 | 88.91 | 11.28 | 0.67 | 1.43 | 0.17 | 37.70 | 54.73 | 15.49 | 52.47 |
| 175.20 | 547.54 | 35.11 | 1.51 | 1.83 | 0.35 | 98.46 | 99.38 | 41.61 | 114.01 |

| Nb (mg/kg) | Cd (mg/kg) | Sn (mg/kg) | Sb (mg/kg) | Cs (mg/kg) | Ba (mg/kg) | La (mg/kg) | Ce (mg/kg) | Pr (mg/kg) | Nd (mg/kg) |
|---|---|---|---|---|---|---|---|---|---|
| 7.92 | 0.71 | 2.14 | 0.82 | 5.87 | 641.33 | 28.34 | 47.82 | 6.88 | 26.11 |
| 19.67 | 0.02 | 4.29 | 0.93 | 12.24 | 337.19 | 24.19 | 71.51 | 4.34 | 14.03 |
| 7.00 | 0.39 | 1.93 | 0.84 | 6.47 | 507.67 | 27.90 | 44.35 | 7.22 | 27.46 |
| 11.95 | 0.50 | 3.22 | 0.92 | 9.77 | 510.60 | 31.45 | 59.02 | 6.98 | 24.96 |
| 7.13 | 0.02 | 1.37 | 0.42 | 5.32 | 334.08 | 19.69 | 43.33 | 4.54 | 16.53 |
| 9.08 | 0.27 | 1.72 | 0.55 | 7.08 | 329.92 | 23.72 | 48.48 | 5.70 | 20.22 |
| 8.08 | 0.55 | 1.89 | 0.51 | 6.31 | 391.84 | 22.08 | 47.22 | 5.11 | 18.40 |
| 11.09 | 0.85 | 3.10 | 0.88 | 7.46 | 456.59 | 28.44 | 58.04 | 6.46 | 23.27 |
| 16.74 | 0.13 | 3.45 | 0.90 | 11.34 | 354.34 | 44.97 | 86.14 | 10.68 | 37.55 |
| 5.84 | 0.70 | 1.81 | 0.58 | 5.80 | 417.65 | 21.50 | 39.17 | 5.14 | 19.04 |
| 14.20 | 0.12 | 2.70 | 0.76 | 10.48 | 430.99 | 33.30 | 74.64 | 7.06 | 24.27 |
| 2.17 | 0.68 | 1.07 | 0.39 | 3.47 | 413.42 | 16.80 | 28.94 | 4.60 | 18.14 |
| 2.57 | 0.89 | 1.36 | 0.41 | 4.58 | 489.51 | 19.30 | 33.96 | 5.45 | 22.19 |
| 11.32 | 0.73 | 2.45 | 0.82 | 10.07 | 522.15 | 33.86 | 63.28 | 7.87 | 28.43 |
| 15.34 | 0.36 | 3.04 | 0.89 | 12.93 | 607.20 | 43.67 | 82.87 | 9.92 | 34.13 |
| 12.77 | 0.53 | 3.20 | 0.96 | 10.36 | 593.44 | 34.40 | 65.01 | 7.30 | 25.82 |
| 10.22 | 0.59 | 2.98 | 0.91 | 10.50 | 568.51 | 36.88 | 66.56 | 8.49 | 31.28 |
| 9.31 | 0.81 | 2.89 | 1.00 | 9.56 | 599.11 | 33.80 | 65.28 | 8.11 | 30.11 |
| 10.70 | 0.84 | 3.12 | 1.02 | 10.73 | 625.84 | 37.52 | 67.56 | 8.71 | 31.87 |
| 9.90 | 0.65 | 3.22 | 0.95 | 10.85 | 584.79 | 37.74 | 66.13 | 8.65 | 31.69 |
| 10.90 | 0.62 | 3.30 | 0.98 | 11.06 | 586.86 | 37.18 | 66.06 | 8.39 | 30.72 |
| 9.87 | 0.52 | 3.02 | 0.87 | 10.10 | 514.46 | 32.97 | 62.22 | 7.53 | 27.34 |
| 11.94 | 0.65 | 3.51 | 1.06 | 11.86 | 578.36 | 38.12 | 71.22 | 8.36 | 30.26 |
| 10.54 | 0.20 | 1.33 | 0.35 | 5.91 | 220.34 | 20.29 | 40.00 | 4.86 | 17.05 |
| 9.33 | 0.18 | 1.62 | 0.48 | 7.42 | 231.23 | 25.39 | 49.63 | 6.18 | 22.15 |
| 0.09 | 0.26 | 0.17 | 0.25 | 0.27 | 65.70 | 4.80 | 8.18 | 1.52 | 6.23 |
| 5.54 | 0.38 | 1.50 | 0.57 | 4.55 | 408.45 | 18.98 | 32.47 | 4.60 | 16.85 |
| 5.05 | 0.58 | 1.55 | 0.44 | 3.18 | 376.96 | 15.34 | 29.67 | 3.67 | 13.93 |
| 10.11 | 1.92 | 2.33 | 0.87 | 10.23 | 1227.82 | 41.07 | 74.86 | 9.12 | 32.98 |

| Sm (mg/kg) | Eu (mg/kg) | Gd (mg/kg) | Tb (mg/kg) | Dy (mg/kg) | Ho (mg/kg) | Er (mg/kg) | Tm (mg/kg) | Yb (mg/kg) | Lu (mg/kg) |
|---|---|---|---|---|---|---|---|---|---|
| 5.52 | 1.43 | 5.13 | 0.74 | 4.68 | 0.95 | 2.57 | 0.36 | 2.37 | 0.34 |
| 2.79 | 0.58 | 2.56 | 0.47 | 3.06 | 0.66 | 2.04 | 0.31 | 2.28 | 0.34 |
| 5.84 | 1.45 | 5.64 | 0.85 | 5.21 | 1.11 | 3.06 | 0.42 | 2.83 | 0.41 |
| 5.18 | 1.26 | 4.94 | 0.76 | 4.45 | 0.95 | 2.70 | 0.38 | 2.61 | 0.40 |
| 3.33 | 0.75 | 3.03 | 0.46 | 2.77 | 0.58 | 1.57 | 0.22 | 1.56 | 0.23 |
| 3.98 | 0.96 | 3.86 | 0.59 | 3.59 | 0.74 | 2.07 | 0.28 | 2.01 | 0.30 |
| 3.77 | 0.88 | 3.44 | 0.53 | 3.35 | 0.67 | 1.86 | 0.28 | 1.85 | 0.26 |
| 4.83 | 1.06 | 4.12 | 0.64 | 3.79 | 0.77 | 2.21 | 0.33 | 2.18 | 0.32 |
| 7.36 | 1.55 | 5.74 | 0.87 | 5.07 | 0.96 | 2.77 | 0.38 | 2.73 | 0.41 |
| 3.97 | 0.97 | 3.69 | 0.54 | 3.40 | 0.71 | 2.00 | 0.28 | 1.91 | 0.27 |
| 4.54 | 1.00 | 3.80 | 0.58 | 3.59 | 0.70 | 2.13 | 0.31 | 2.09 | 0.32 |
| 3.86 | 0.99 | 3.84 | 0.56 | 3.48 | 0.75 | 2.06 | 0.29 | 1.83 | 0.29 |
| 4.57 | 1.29 | 4.75 | 0.71 | 4.36 | 0.91 | 2.43 | 0.33 | 2.26 | 0.32 |
| 5.60 | 1.38 | 4.98 | 0.78 | 4.57 | 0.95 | 2.61 | 0.36 | 2.43 | 0.37 |
| 6.66 | 1.54 | 5.75 | 0.85 | 5.23 | 1.06 | 2.87 | 0.40 | 2.80 | 0.41 |
| 5.32 | 1.30 | 4.97 | 0.76 | 4.92 | 1.03 | 2.88 | 0.43 | 2.89 | 0.42 |
| 6.47 | 1.54 | 6.11 | 0.93 | 5.77 | 1.20 | 3.37 | 0.47 | 3.17 | 0.46 |
| 6.80 | 1.55 | 6.00 | 0.94 | 5.62 | 1.16 | 3.32 | 0.49 | 3.11 | 0.46 |
| 6.61 | 1.63 | 6.19 | 0.93 | 5.66 | 1.21 | 3.26 | 0.47 | 3.11 | 0.47 |
| 6.50 | 1.60 | 6.19 | 0.93 | 5.53 | 1.18 | 3.28 | 0.47 | 3.08 | 0.45 |
| 6.40 | 1.48 | 5.92 | 0.90 | 5.49 | 1.14 | 3.22 | 0.46 | 3.09 | 0.45 |
| 5.58 | 1.34 | 5.25 | 0.80 | 5.01 | 1.02 | 2.87 | 0.41 | 2.75 | 0.41 |
| 6.14 | 1.42 | 5.48 | 0.86 | 5.20 | 1.07 | 2.98 | 0.43 | 2.89 | 0.42 |
| 3.29 | 0.77 | 2.88 | 0.44 | 2.47 | 0.49 | 1.36 | 0.20 | 1.34 | 0.19 |
| 4.45 | 1.02 | 3.80 | 0.57 | 3.34 | 0.66 | 1.71 | 0.26 | 1.77 | 0.25 |
| 1.64 | 0.49 | 1.69 | 0.25 | 1.42 | 0.27 | 0.60 | 0.07 | 0.48 | 0.06 |
| 3.52 | 0.87 | 3.20 | 0.49 | 2.96 | 0.59 | 1.66 | 0.23 | 1.55 | 0.23 |
| 3.03 | 0.78 | 2.79 | 0.43 | 2.57 | 0.55 | 1.52 | 0.21 | 1.43 | 0.20 |
| 6.69 | 1.63 | 6.28 | 0.96 | 5.87 | 1.25 | 3.53 | 0.49 | 3.32 | 0.48 |

| Hf (mg/kg) | Ta (mg/kg) | W (mg/kg) | Tl (mg/kg) | Pb (mg/kg) | Bi (mg/kg) | Th (mg/kg) | U (mg/kg) |
|---|---|---|---|---|---|---|---|
| 2.42 | 0.61 | 0.32 | 0.60 | 12.29 | 0.19 | 9.28 | 1.83 |
| 5.04 | 1.58 | 3.59 | 0.84 | 16.78 | 0.53 | 18.63 | 2.65 |
| 2.28 | 0.53 | 0.27 | 0.53 | 12.36 | 0.23 | 8.66 | 1.58 |
| 3.55 | 0.94 | 0.16 | 0.63 | 19.21 | 0.36 | 15.05 | 2.48 |
| 2.32 | 0.51 | 1.16 | 0.45 | 14.75 | 0.19 | 6.26 | 1.41 |
| 2.77 | 0.65 | 1.50 | 0.52 | 15.50 | 0.27 | 8.18 | 1.47 |
| 2.58 | 0.61 | 1.69 | 0.62 | 16.36 | 0.50 | 8.05 | 1.54 |
| 3.30 | 0.90 | 0.14 | 0.60 | 24.20 | 0.53 | 14.10 | 2.54 |
| 4.64 | 1.30 | 2.35 | 0.78 | 23.37 | 0.41 | 16.98 | 2.28 |
| 2.13 | 0.42 | 0.20 | 0.53 | 2.89 | 0.16 | 8.00 | 1.46 |
| 4.10 | 1.07 | 1.23 | 0.60 | 19.91 | 0.28 | 13.79 | 1.79 |
| 1.20 | 0.13 | 0.06 | 0.36 | 1.07 | 0.07 | 4.99 | 1.16 |
| 1.52 | 0.16 | 0.09 | 0.42 | 2.26 | 0.17 | 5.97 | 1.39 |
| 3.27 | 0.84 | 0.48 | 0.65 | 18.43 | 0.40 | 12.33 | 1.78 |
| 4.22 | 1.10 | 0.40 | 0.79 | 22.81 | 0.35 | 15.36 | 2.19 |
| 4.06 | 1.00 | 0.38 | 0.77 | 18.59 | 0.32 | 15.76 | 2.59 |
| 3.34 | 0.75 | 0.15 | 0.60 | 11.81 | 0.27 | 13.04 | 2.16 |
| 3.46 | 0.66 | 0.08 | 0.55 | 10.48 | 0.26 | 11.94 | 2.12 |
| 3.52 | 0.78 | 0.10 | 0.72 | 9.22 | 0.22 | 13.42 | 2.06 |
| 3.51 | 0.68 | 0.22 | 0.59 | 9.54 | 0.18 | 13.60 | 2.23 |
| 3.56 | 0.79 | 0.17 | 0.70 | 10.90 | 0.27 | 13.93 | 2.19 |
| 3.38 | 0.73 | 0.09 | 0.63 | 11.70 | 0.28 | 13.30 | 2.08 |
| 3.86 | 0.86 | 0.36 | 0.74 | 11.09 | 0.26 | 15.33 | 2.26 |
| 2.17 | 0.60 | 0.61 | 0.36 | 12.18 | 0.19 | 7.34 | 1.16 |
| 2.56 | 0.67 | 0.47 | 0.45 | 13.64 | 0.18 | 8.74 | 1.40 |
| 0.02 | 0.01 | 0.00 | 0.03 | 1.33 | 0.24 | 0.25 | 0.40 |
| 1.63 | 0.45 | 0.39 | 0.35 | 9.89 | 0.22 | 6.37 | 1.21 |
| 1.56 | 0.39 | 0.17 | 0.34 | 8.25 | 0.14 | 5.54 | 1.30 |
| 3.37 | 0.81 | 0.03 | 0.48 | 13.03 | 0.26 | 12.89 | 2.07 |
