## Supplementary Information for "Evidence for deliberate burial of the dead by *Homo naledi*"

### Supplementary Information 1: Description of the sedimentary deposits of the Dinaledi Subsystem

The sedimentary deposits within the Dinaledi Chamber and Hill Antechamber have been categorized within three units defined on lithological but not necessarily chronological bases (Dirks et al. 2015; Dirks et al. 2017; Wiersma et al. 2019). An initial description was provided by Dirks et al. (2015) and this description was revised by Dirks et al. (2017); Wiersma et al. (2019) presented additional data and a hypothesis for the processes of autobrecciation of sediment deposits within the subsystem. This section reviews these previous descriptions so that readers of the current study have the relevant sediment descriptions available.

**Unit 1** consists of unlithified, horizontally laminated orange-red mudstone, abbreviated as LORM. Deposits of this unit formed when water laden with mud filtered into the cave chamber and deposited material that was suspended within it. The content of Unit 1 facies varies in grain size and composition, with some sandy lenses and inclusion of microfaunal remains. Outcrops of Unit 1 are localized within fissure passages extending from the chamber and on chert ledges on chamber walls. Unit 1 includes the oldest sedimentary deposits in the chamber as inferred from the presence of Unit 1 clasts within Unit 2 and Unit 3 sediments. But outcrops of Unit 1 on chert ledges and in fissure passages were deposited later and are still being deposited today, so that this unit is time-transgressive relative to the other sedimentary units in the subsystem (Dirks et al. 2017; Wiersma et al. 2019; see supplementary figure S1).

Unit 2 and Unit 3 both include clasts of Unit 1 laminated orange-red mudstone (LORM) embedded within a brown clay matrix. These clasts are angular or subangular, they vary in size but are typically less than 3 cm in diameter, and they do not retain any orientation from their original deposition, with the originally-horizontal laminations now at random orientations.

**Unit 2** includes dolomite and chert fragments and is found only beneath a flowstone sheet near the entrance of the Hill Antechamber (Dirks et al. 2017); the oldest part of this flowstone sheet (Flowstone 1a) has reversed magnetic polarity and is older than 780 ka, which is a minimum age for these Unit 2 sediments.

**Unit 3** outcrops across the remainder of the floor of both the Hill Antechamber and Dinaledi Chamber. We have distinguished two sub-units of Unit 3: the lower **sub-unit 3a** which lacks *H. naledi* fossil material and upper **sub-unit 3b** which contains *H. naledi* fossils. The earliest known incidence of *H. naledi* in sub-unit 3b sediments is constrained at less than 335 ka by direct dating of *H. naledi* teeth (Dirks et al., 2017). A direct-dated baboon tooth 35 to 40 cm below *H. naledi* material suggests a maximum age for this part of sub-unit 3a of 723 ± 181 ka (Dirks et al. 2017). These dates demonstrate that the formation of Unit 2 and Unit 3 sediments was neither rapid nor simultaneous with the arrival and emplacement of *H. naledi* remains in sub-unit 3b sediments.

We have previously reported on the composition and micromorphology of sub-unit 3b sediments from two areas of the Dinaledi Chamber. This work included micromorphological and bulk sediment samples from a surface surrounding *H. naledi* remains and from within the 2013 excavation unit directly in contact with subsurface *H. naledi* remains (Dirks et al. 2015). We summarize these results briefly here. These samples exhibit the same grain size and shape distributions, predominantly composed of clay fragments with angular shapes, in random orientations without evident bias or preference to horizontal. The micromorphology and composition evident in these samples reflect the formation history of sub-unit 3b deposits that are in direct contact with *H. naledi* remains. These samples contrast strongly with sediments of the nearby Dragon's Back Chamber, which have a high input of detrital quartz and stony clasts from outside the cave system. Unlike the Dragon's Back Chamber sediments, sub-unit 3b sediments in the Dinaledi Chamber have little quartz content and are dominated by clay minerals and mica, with a texture composed of angular clay fragments and laminated orange-red mud (LORM) clasts (Dirks et al. 2015; Wiersma et al. 2019). These LORM clasts are remnants of Unit 1 sediments that were reworked during the formation of sub-unit 3a sediments. The laminations within LORM clasts in sub-unit 3b are oriented randomly, and this orientation reflects the origin of sub-unit 3b from weathering and reworking of Unit 1 and sub-unit 3a sediments. The micromorphology of sub-unit 3b samples demonstrates that the formation of this sub-unit occurred under low energy conditions with minimal lateral transport of material and without sufficient water saturation to dissolve LORM clasts or to remove sharp angles from them (supplementary figure S2).

**Mud.** Previous work has employed the geological terms "mud", "mudstone", and "mud-clasts" in the description of sediments within the Dinaledi Subsystem (Dirks et al. 2015; Dirks et al. 2017; Wiersma et al. 2019). These terms have been misunderstood by some commentators, who have assumed that the term "mud" must imply that water actively flowed through the Dinaledi Chamber at the time of deposition of *H. naledi* remains. This misconception about the nature of the Dinaledi Chamber deposits can be avoided by differentiating the different units and their formation history.

The laminations in Unit 1 sediments formed when water carried fine sediment into the subsystem, some of which percolated through cracks and deposited material on chert ledges and some in fissure passages. These laminations resulted from successive introductions of water-saturated sediments onto surfaces; at the time of their formation the laminations were horizontal or parallel to underlying surfaces such as chert ledges (Wiersma et al. 2019). Unit 1 sediments remain in these places today. No *H. naledi* or other macrofaunal fossil material has been found in Unit 1 sediments. All *H. naledi* material occurs within Unit 3 sediments, interpreted as having formed by autobrecciation through the weathering and reworking of Unit 1 (Dirks et al. 2015; Wiersma et al. 2019). Where we excavated Unit 3 deposits and where they are exposed on the surface, they consist of an unconsolidated, loosely-packed clay matrix containing clasts and frequent air voids. We have divided Unit 3 into two sub-units: sub-unit 3b contains *H. naledi* remains while sub-unit 3a lacks them. Both contain clasts derived from Unit 1, and these clasts are angular in outline and retain their laminations, but the laminations in different clasts are not aligned with each other or with a horizontal plane. In micromorphological thin sections, the LORM clasts appear at random orientations (Dirks et al. 2015). The micromorphology of sub-unit 3b sediments is inconsistent with flowing water, high-energy transport, or saturation sufficient to round the soft angular clasts or dissolve them. Sub-unit 3b sediments contain moisture: the air humidity within the cave averages 97%–99%, there are drip points at various places with some speleothem deposition, and the *H. naledi* skeletal material contains moisture and is fragile. The use of the term "mud" does not denote a formation history with flowing water, and the sedimentology of sub-unit 3b sediments rules out flowing water as a cause for their formation (supplementary figure S3).

**Reworking and autobrecciation.** Previous work (Dirks et al. 2015; Wiersma et al. 2019) discusses the reworking and autobrecciation of sediments in the Dinaledi subsystem. The mud-clast breccia of Unit 2 and sub-unit 3a in the Dinaledi Subsystem formed as a result of weathering, erosional breakdown, and reworking of Unit 1 sediments, resulting in unlithified breccia units containing clasts derived from Unit 1 sediments. This breakdown was a result of several processes. The composition of sediments includes clay minerals that are susceptible to swelling and contraction with greater or lesser moisture content. Fluctuations in moisture in the cave system such as those accompanying seasonal or longer cycles in the environment external to the cave resulted in the relative desiccation of parts of Unit 1 sediments and the formation of fine cracks (Wiersma et al. 2019). Such fine cracks in some areas led to infiltration of the Unit 1 sediments by calcite and aragonite, which is visible in some parts of the cave system today. Microbial processes and oxidation of iron and manganese also were part of this diagenesis of Unit 1 sediments. The area of the Hill Antechamber exhibits a substantial (~ 30 degree) floor slope today, and cycles of humidity with expansion and contraction might lead to some horizontal displacement of sediments on such a slope (Dirks et al. 2015; Wiersma et al. 2019).

In previous work, Dirks and coworkers (2015) suggested that such cycles of humidity may have continued after the deposition of *H. naledi* remains and may have contributed to the fragmentation of *H. naledi* skeletal material, explaining in part the post-depositional fracturing of the material. In that work it was noted that some skeletal elements were stained by mineral precipitates forming "tide marks" that are recognizable on anatomically contiguous fragments after refitting. Some of these elements were further fragmented after the tide marks formed, and some of those fragments shifted in position. In addition, some skeletal elements were uncovered in positions that could not have occurred on a horizontal surface or within a shallow depression, including near-vertical orientations of long bone shafts (Dirks et al., 2015).

The evidence of post-depositional fragmentation and non-horizontal orientations appeared to contrast with other elements and skeletal regions that were deposited as whole limbs retaining articulations of elements. These aspects of the deposit are not consistent with the deposition of bone on a near-horizontal floor surface followed by slow sedimentation. As we noted, these spatial properties of the sediment and bone must have resulted from some kind of highly selective reworking of the sub-unit 3b (Dirks et al. 2015; Dirks et al. 2016). Some *H. naledi* bones show evidence of invertebrate activity, and previous work has suggested the possibility of some bioturbation by these agents in the sediments (Wiersma et al. 2019).

The excavations that we report here have given us some additional information about the nature of reworking in Unit 3 sediments. Our excavation units in the Hill Antechamber include areas with a higher density of Unit 1-derived clasts, enough to form rough layers parallel to the sloping floor surface. These layers are interrupted by the intrusion of skeletal material. In the Dinaledi Chamber, our excavation encountered one horizontal layer rich in clasts that is interrupted by a feature containing skeletal remains.

In our previous considerations we neglected two possible causes of reworking in our previous considerations of the Dinaledi Subsystem that could result in the patterns we observed: reworking of sediment due to collapse of spaces left by decomposition of soft tissue, and reworking by *H. naledi*. In the current study we consider these processes for the first time.

#### Supplementary Information 2: Entry of *Homo naledi* into the cave system

##### Supplementary Information 2.1: Entrances to the cave system

The contiguous areas of the Rising Star cave system that are accessible today connect to the surface outside the cave via several entrances along the east and northeast-facing hillside above the system. The Dinaledi Subsystem is approximately 30 m beneath the present surface, and the nearest opening to the outside is via the Dragon's Back Chamber and Postbox Chamber, a horizontal distance of more than 100 m and vertical distance of 30 m (Dirks et al. 2015). Team members who work in the cave today usually follow a somewhat longer and easier route into the Dragon's Back Chamber via the Superman Chamber. Both these routes into the system involve vertical descents, sharp turns, and squeezes through rock passages of less than 30 cm.

The southwestern-most extent of the Dragon's Back Chamber is tall, narrow and largely filled by a high, jagged fin of rock, approximately 10 m in length and 7 m in height, called the Dragon's Back. This structure is a dolomite septum that separates narrow spaces on either side, which become too narrow at the floor level to accommodate passage, and which terminate before the chamber's southwestern end. The chert banding of the Dragon's Back itself is displaced approximately 60 cm downward relative to the bedrock of the surrounding chamber walls, indicating a shift in the position of this structure sometime in the past (Dirks et al. 2015; Robbins et al. 2021). Today it is possible to climb the Dragon's Back structure and then descend 12 m through a fissure network at its southwestern extent, which passes into the Dinaledi subsystem. This fissure network opens from above at several places along the top of the Dragon's Back. In 2013 Steven Tucker and Rick Hunter descended the most distal of these passages for the first time and recognized skeletal remains in the Dinaledi subsystem, leading to the exploration of this space and recovery of *H. naledi* (Berger et al. 2015; Dirks et al. 2015). This passage from Dragon's Back to the Hill Antechamber became known as the Chute, and it cannot be traversed by humans who cannot fit their bodies through a squeeze of less than 18 cm. There are other pathways through this fissure network, but exploration of them did not identify any passage that could accommodate any larger person (Elliott et al. 2021). One of these pathways is used today for cables that support our work in the Dinaledi subsystem.

The Lesedi Chamber is in a different part of the cave system. There is no path directly from the Lesedi Chamber to the Dinaledi subsystem; the closest route is more than 130 m. The Lesedi Chamber is approximately 30 m below surface and 86 m from the closest current outside surface entrance. This chamber can be entered from three different directions and all of these involve vertical descents, tight turns, and squeezes of 30 cm or less.

The complexity of accessing these spaces today has brought a great deal of attention to the question of whether *Homo naledi* entered the cave system and these chambers by the same route as today's scientists. Cave systems change over time and the Rising Star system is no exception. The time involved for these changes since the occurrence of *H. naledi* is more limited than for some nearby cave systems with remains of Pliocene and Early Pleistocene species such as *Australopithecus africanus* or *Paranthropus robustus*. Present evidence shows that some *H. naledi* remains entered the Dinaledi Subsystem between 335,000 and 241,000 years ago (Dirks et al. 2017; Robbins et al. 2021). No geochronological estimate yet exists for the Lesedi Chamber material, and the Dinaledi Subsystem results do not exclude the possibility that some individuals may represent a broader time range.

Most cave systems in the region were affected by lime mining activity during the early twentieth century. In the Rising Star cave system, the southern entrance of the cave and adjacent underground areas were affected the most strongly; neither of these areas is closer than 250 m through the cave system from the Dragon's Back Chamber. The eastern entrance to the system passes through the Skylight Chamber, and miners here blasted a small tunnel and constructed a ramp to gain easier access into this chamber. There is no visible trace of mining activity in the Dragon's Back Chamber or Postbox Chamber, or in the areas adjacent to the Lesedi Chamber.

Our investigations of the cave system, flowstone geochronology, and sedimentation processes have helped build a partial understanding of how the system changed in the period between 600,000 years ago and recent times (Dirks et al. 2015; Dirks et al. 2017; Wiersma et al. 2019; Elliott et al. 2021; Robbins et al. 2021). Detrital quartz and other material derived from surface contexts external to the cave system is abundant within lithified breccia deposits in the Dragon's Back Chamber and unlithified sediment deposits that occur in the Postbox Chamber and Superman Chamber. The current nearest entrances to the system, such as the Skylight Chamber, are not situated in a way that sediment input from them could fill large areas of the Postbox, Superman, or Dragon's Back chambers from these sources. Survey of the upper areas of the Postbox Chamber suggest a possible scenario for the entrance of this external material into the system, described in Robbins et al. (2021). Much of the externally derived sediment is in positions where gravity could carry material from the highest parts of the Postbox Chamber, which are at the northeastern-most extent of the chamber. This area is below the point where two large chert horizons outcrop on the hillside above the Skylight Chamber. If an entrance once existed at this location, it must now be completely filled with sediment and is not obvious either from the surface or from within the Postbox Chamber. Any entrance at this location would have opened to a vertical descent of approximately 10 m into the upper part of the Postbox Chamber. From such an entrance, the distance from the surface to the Dinaledi Subsystem would be reduced by up to 20 m, making it around 80 m in total.

No stones or large grains of externally derived sediment input occur in the Dinaledi Subsystem sediments (Dirks et al. 2015; Wiersma et al. 2019). This indicates that no passive gravity-driven route for sediment input from the surface into this subsystem existed throughout the formation of Units 1, 2, or 3, inferred to be the last 700,000 years or more (Dirks et al. 2017).

A 5 meter sill occurs within the Dragon's Back Chamber, separating the main area of the chamber from the southwestern section in which the Dragon's Back ridge is situated (Robbins et al. 2021). This sill prevented direct sediment flow from the main area of the Dragon's Back Chamber up to the floor near the Dragon's Back itself, and thereby closed off sediment flow from there toward the Dinaledi Subsystem (Robbins et al. 2021). Sediment input into the Dinaledi Subsystem has occurred by slow introduction of fine sediment with dripwater at various points in the subsystem, as well as slow weathering of dolomite. The sediment floor level is highest at the northeasternmost extent of the Hill Antechamber and this may suggest some input of material from the fissure network that separates this subsystem from the Dragon's Back Chamber. However, neither the sediment thickness nor the underlying bedrock floor elevation is known within the Hill Antechamber. The 30 degree floor slope is higher than the 15 degree southwest-trending dip of the dolomite bedrock, but may not indicate a very great accumulation of sediment in this area.

The accessibility of the Dinaledi Subsystem may have been altered over time by changes to the Dragon's Back structure itself. Robbins et al. (2021) developed a chronology of the deposition of massive orange sand (MOS) and laminated orange sand (LOS) breccias within the Dragon's Back Chamber, establishing that the MOS breccia was formed between 290 ka and 225 ka. The MOS breccia contains large dolomite blocks, and these authors hypothesize that the formation of this breccia may correspond to a period of changes to the roof and structure of this part of the system. They further suggest that it is a reasonable hypothesis that the Dragon's Back structure itself may have taken its current form at that time. The 60 cm downward displacement and 10 degree rotation of this large block would have changed the accessibility of this space within the Dragon's Back chamber. Robbins et al. (2021) suggest that passage beneath this block may have been possible prior to its displacement.

However this displaced block alone does not form the separation between the Dragon's Back Chamber and the Hill Antechamber. These areas were connected prior to the block displacement as they are today: only through various small passages of a fissure network in the dolomite bedrock. The displaced block does not define the vertical descent that today provides the most common access into the Dinaledi Subsystem via this fissure. Passage beneath the block prior to its displacement may have provided greater access into this fissure network but the extent to which this may have affected ingress into the Dinaledi Subsystem is not evident. Work to characterize the constraints of the present very challenging passageways continues.

In summary, *H. naledi* localities all are located a minimum of 86 meters from the nearest identifiable current entrance into the cave system, and one hypothesized past entrance may reduce this minimum to approximately 80 m. All localities are accessible only through narrow passages that include vertical descents of at least 30 m, sharp turns, and squeezes of less than 25 cm, and remains the case for any hypothesized past entrance. While many commentators have emphasized the physical difficulty of these passages for humans today, most aspects of the anatomy of *H. naledi* appear to be well-suited for movement within these spaces. With adult body mass of less than 50 kg, stature less than 160 cm, thin and long lower limbs, shoulders with a cranially oriented glenoid fossa, strong and long thumb, and a powerful grip, the locomotor and body size adaptations of *H. naledi* are matched to climbing and moving into passages with a small diameter.

**Supplementary Information 2.2: Hypotheses tested in previous work**

In previous work, we have reported some ways that the hominin fossil material in the Dinaledi Chamber, Lesedi Chamber, and U.W. 110 locality depart from the situation in other fossil-bearing sites in the Cradle of Humankind, and we have addressed several hypotheses related to the deposition of these remains (Dirks et al. 2015; Dirks et al. 2016; Hawks et al. 2017; Elliott et al. 2021; Brophy et al. 2021). Remains of *H. naledi* are situated within chambers and spaces at substantial distances from each other, and at substantial distances from current entrances into the system. These localities have a limited representation of non-hominin faunal material and a high abundance of *H. naledi* with at least 16 individuals represented in the Dinaledi Subsystem prior to the current study, and at least 3 individuals in the Lesedi Chamber. Our initial analysis of both the Dinaledi Chamber and Lesedi Chamber situations documented some intact articulations in the skeletal remains from these localities (Berger et al. 2015; Dirks et al. 2015; Hawks et al. 2017). Subsequent work showed spatial clustering of elements within the Dinaledi Chamber subsurface excavation (Kruger et al. 2016) and correlated one of these clusters with elements from a single subadult partial skeleton (Bolter et al. 2020). The retention of anatomical articulations and clustering of material from particular individuals demonstrated that some skeletal material was emplaced in sediment within the Dinaledi Chamber and within the Lesedi Chamber prior to the decomposition of soft tissue (Dirks et al. 2015; Hawks et al. 2017). Excavation uncovered some skeletal elements of *H. naledi* in subvertical orientations while in close contact with other elements in near-horizontal or orientations, a pattern inconsistent with the deposition of bones on a horizontal surface or in a shallow depression. We interpreted this pattern as suggesting multiple depositional events (Dirks et al. 2015).

In our past work, we have tested several hypotheses for the presence of hominin remains in various parts of the Rising Star cave system. Here we accept the results of this previous work and review the evidence that rejects several specific hypotheses.

**Carnivore accumulation.** At a few other cave sites in South Africa, large carnivores interacted with some hominin remains, as evidenced by tooth scoring, punctures, and other carnivore damage on some hominin fossil bones (Brain, 1985 & 1993). Some researchers have proposed that the remains of *H. naledi* from the Dinaledi Chamber may have been accumulated by large carnivores, pointing either to the pattern of fragmentation of the remains or the skeletal part representation (Val 2016; Egeland et al. 2018).

The hypothesis of carnivore accumulation is testable with reference to the remains and their context. We have previously shown that no element from either the Dinaledi Subsystem or Lesedi Chamber have any tooth scoring or punctugoffres, and none have breakage consistent with carnivore activity on fresh bone (Dirks et al. 2015; Hawks et al. 2017; Brophy et al. 2021). All fractures evident in the Dinaledi Chamber and Lesedi Chamber fossil material are consistent with breakage after deposition, burial, and subsequent loss of organic content (Dirks et al. 2015; Hawks et al. 2017). Additionally, non-hominin macrofaunal elements are rare and do not include other potential large prey animals (Dirks et al. 2015; Hawks et al. 2017). Excavations and surface examination of both areas has failed to yield any evidence of carnivore presence within them. There are no coprolites, no evidence of burrowing or other sediment disruption by large carnivores, and no large carnivore remains. In contrast to the Dinaledi Subsystem, the Lesedi Chamber assemblage does present evidence of small carnivores, including mongoose and small canids (Hawks et al. 2017), but such species have not been implicated in predation of hominins and cannot have transported or accumulated hominin bodies within a deep cave. These observations falsify the hypothesis that carnivores played any role in the presence of hominins in these parts of the cave system.

**Passive gravity-driven accumulation.** Dirks and coworkers (2015) discussed the hypothesis that hominin remains within the Dinaledi Chamber may have been introduced at or near our current access point, dropping into the Hill Antechamber and spreading from there under the influence of gravity alone. This scenario would resemble that proposed for the entrance of hominin remains into the Sima de los Huesos, Spain (reviewed by Aranburu et al. 2017). The evidence from previous work (Dirks et al. 2015) was sufficient to raise doubts about this hypothesis for several reasons. Remains are found in the Dinaledi Chamber, more than 20 m from the nearest access point into the subsystem, and separated from the Hill Antechamber by a long (> 7 m) and narrow (< 0.5 m) fissure passage. This fissure passage includes the lowest point in the subsystem, adjacent to drains that could channel sediment downward (Dirks et al. 2015). Articulated bodies and parts of bodies within the Dinaledi Chamber could not have traversed this route under the force of gravity alone. Our subsequent work in the subsystem produced further evidence refuting this hypothesis. The presence of hominin remains in the fissure network distal to the Dinaledi Chamber, including the U.W. 110 locality, cannot have arrived in their present locations passively by gravity alone (Elliott et al. 2021; Brophy et al. 2021). This hypothesis for the Dinaledi Subsystem is further refuted by the results of the current study. The additional excavation units within the Dinaledi Chamber and Hill Antechamber show clearly that hominin remains are concentrated in specific features and do not form a talus or continuous layer in either the Dinaledi Chamber or Hill Antechamber. In addition, the hominin material within the Lesedi

Chamber is within a different depositional space far from the Dinaledi subsystem, with different access constraints that are not consistent with passive, gravity-driven introduction of hominin remains.

**Occupation.** The long-term use of the Dinaledi Subsystem by *H. naledi* for occupation appears to be inconsistent with the fossil and sedimentary situation. No occupation debris or other evidence of occupation has been identified within this subsystem or within the Lesedi Chamber.

**Water transport and burial by fluvial action.** As discussed above, the sediments of sub-unit 3b within the Dinaledi Subsystem did not form in flowing water or with saturation sufficient to round the shape of LORM clasts and angular clay fragments which comprise this sub-unit (Dirks et al. 2015; Wiersma et al. 2019). These micromorphological observations are derived from sediment found in direct contact with surface and subsurface fossil remains of *H. naledi* (Dirks et al. 2015). Additionally, water transport of either bodies, large parts of bodies, or bones is refuted by physical constraints of the system and evidence from the surfaces of bone elements themselves (Dirks et al. 2015; Dirks et al. 2016). The Dinaledi Subsystem is separated from the adjacent Dragon's Back Chamber by a ~5 m sill of dolomite that has been in place throughout the time of formation of Unit 3 (Robbins et al. 2021). This has prevented the movement of sediment directly into the Dinaledi Subsystem from Dragon's Back or other areas nearer to surface entrances, and explains the sedimentological contrasts between the two chambers (Dirks et al. 2015). This leaves no possibility for bodies or bones to have been transported into Dinaledi by water flow. The transport of bodies by water from the Hill

Antechamber into the Dinaledi Chamber is likewise not possible; bodies would have to flow down one of two narrow passages, past floor drains that would channel any water further down into the cave, and after passing through these passages would then have had to continue with minimal grade or even uphill for at least 15 meters to their resting place in the Dinaledi Chamber. The transport of bodies or bones by water into the distal fissure network is likewise not credible (Brophy et al. 2021). Water transport of bodies into the Lesedi Chamber has not been suggested and cannot explain the distribution of skeletal material in the chamber.

**Mass mortality or death trap.** Our initial exploration of the Dinaledi Chamber produced evidence of a minimum number of 15 individuals representing ages from infant to old adult (Berger et al. 2015; Bolter et al. 2018). The distribution of ontogenetic ages of this assemblage could be consistent with either attritional or catastrophic mortality distributions and did not distinguish them (Bolter et al. 2018). These data by themselves appeared to leave open the possibility that some or all of the assemblage might represent individuals that entered the chamber alive and then died there (Dirks et al. 2015; Dirks et al. 2016). Later workers suggested that baboon sleeping sites might provide an analogy for the accumulation of skeletal remains in the Dinaledi Chamber, with bodies gradually accumulating after natural deaths of individuals (Nel 2019; Nel et al. 2021). However, we previously noted several aspects of the Dinaledi hominin assemblage that are inconsistent with an unburied mass death accumulation, including the presence of skeletal elements in subvertical orientations in close spatial proximity or contact with articulated parts of bodies (Dirks et al. 2015; Dirks et al. 2016). The presence of multiple hominin individuals in the Lesedi Chamber and within widely-separated localities in the Dinaledi Subsystem also implicates regular use of extensive parts of the cave system rather than a single mass death event (Hawks et al. 2017; Brophy et al. 2021). The localities with *H. naledi* remains contrast with known baboon sleeping sites in several ways, including their distance from cave entrances, lack of evidence of external input of sediment or organic material, and the extent of bias toward hominin (or primate) remains at the expense of other taxa.

#### Supplementary Information 3: Description of the Hill Antechamber Artefact 1 associated with the Hill Antechamber Feature

Within the Hill Antechamber Feature is a rock of 138.5 mm length which is located within 20 mm of a group of bones interpreted as anatomically associated metacarpals and phalanges of a right hand. Due to the proximity of this rock to the hand remains and its general resemblance to a large flake tool we prioritized obtaining higher-resolution scan data of the stone to evaluate its morphology. We designated it Hill Antechamber Artifact 1 (HAA1) recognizing that at a minimum its spatial association is of interest. At this writing, HAA1 still remains within the larger plaster jacket containing most of the Hill Antechamber feature. This section presents a description of the object. Orientation is described based on an arbitrarily defined position of features illustrated in supplementary figure S21. All measurements and descriptions are taken from 3D images produced from the synchrotron scans or from examination of high-resolution 3D prints of HAA1. Both scans and 3D shape files of HAA1 are available for download on https://Morphosource.org . A high resolution movie of HAA1 is available as Supplementary Movie 2.

The shape of HAA1 is distinctive in comparison to other rocks on the surface or encountered during excavation of the Hill Antechamber or Dinaledi Chamber. HAA1 is 138.5

mm in total length. Its greatest width perpendicular to length is 49mm just left of the middle of center. Its greatest diameter in the remaining dimension is 26.3mm. HAA1 is roughly crescent-shaped coming to a sharp point laterally left and a more rounded but antero posteriorly sharp edge right laterally. It presents a large flake scar of ca. 80mm in length on the anterior-inferior surface from approximately 10mm left laterally of its midline that travels to within 12mm of the right lateral end. The lunate area of this flake scar occupies approximately three quarters of the body of the artifact for most of its extent. A 6-7mm wide ledge is observed in the superior one third of the artifact that is created from the removal of rock flake ca. 84.5mm in length that leaves a prominent hump occupying the middle 65mm of the artifact. We cannot determine the type of stone unambiguously from scan data, but we hypothesize based on its surface characteristics that it is likely composed of dolomite, and there is no evidence that the rock is exogenous to the cave system.

Within the area of the lunate flake scar on the anterior surface, and opposite this area on the posterior surface, striations are visible that appear to be use wear or erosional marks (Figure 12). Some of these lines are as much as 1.5mm wide but most are sub-millimeter in width and are 20 to 25mm in length. They travel predominantly in an lower-left to upper-right direction across this surface (supplementary figure S22).

The posterior surface of HAA1 is dominated by a prominent ridge that runs from the left lateral point superolaterally at an approximate 15-degree angle, reaching the superior edge about 50mm from the right lateral edge. The peak of the ridge forms the greatest width of the artifact and this occurs approximately 55mm from the left lateral end. Supero-inferiorly oriented erosional or wear lines are found on the right lateral half of the artifact roughly mirroring those on the anterior surface. There is a depressed area that may represent a worn flake removal in this same region that is not as large nor prominent as that found on the anterior surface.

The inferior edge of the artifact along the area of the anterior lunate flake scar is very sharp and under high resolution imaging by synchrotron (6µm), irregular serration is visible across the whole of the surface (supplementary figure S22). No other serration is obvious along the edges of other sharp areas.

#### Supplementary Information 4: Hominin skeletal material and element representation in the Dinaledi Feature 1 and 2 area

In this section we present a preliminary identification of skeletal elements within the burial features, with an assessment of skeletal part representation. We note the presence of articulated or spatially contiguous elements where this has been observed.

In both the Dinaledi and Hill Antechamber, we have left much of the evidence described here *in situ* or *en bloc* without unnecessary destructive excavation or preparation. The skeletal material that has been excavated from Dinaledi Features 1 and 2 has not been subjected to solvents or consolidants at this time (Materials and Methods). The Hill Antechamber feature rests within three plaster-jacketed blocks at the present time. These decisions limit the number of observations that we can gather on the skeletal remains from these features. Here it is worth a brief comment to clarify these protocols and discuss their relevance to the study of the burials.

Archaeological controlled excavation is a fundamentally destructive process (Lucas, 2001), one in which the spatial relationships between buried objects (including hominin fossils) is lost once the matrix surrounding such remains is disturbed and the buried objects lifted. Spatial relationships are necessary to determine temporal relationships in archaeological settings; they are also fundamental to understanding the processes of alteration that assemblages may have undergone after burial. Article 5 of the International Charter for Archaeological Heritage Management states that archaeological investigations can be carried out using a wide array of methods from non-destructive remote sensing, through sampling, to total excavation. It is widely recognized that total excavation is a recourse that should be adopted after consideration of other less destructive means of study (Beaudet and Elie, 1991).

3D modeling or non-invasive imaging is the preferred method for recording and understanding fragile, friable buried contexts where the precise spatial relationships between clastic components is considered more important rather than the necessity to view the surface morphology of such remainsi. Such a spatially-based approach has been used and advocated in recording of mainstream archaeology (Roosevelt et al, 2015), cremains (Pankowská et al. 2017), bone taphonomy (Silver, 2016; Randolph-Quinney et al., 2018), 4D documentation of pit burials (Mickleburgh and Westcott, 2018), virtual autopsy (Bolliger and Thali, 2009), recording fragile funerary goods and human remains (Jansen et al., 2006), through to whole burial chambers (Krausse et al, 2017), where information derived from the spatial relationships within the burial environment outweighs that from direct analysis of the bones or artefacts themselves (Dell'Unto and Landeschi, 2022).

In the Rising Star cave system we have employed study protocols that minimize the extent of excavation areas and maximize the collection of spatial data by multiple modalities (Elliott et al 2021; Dirks et al 2015 and 2017; Kruger et al 2016). The total excavation surface in the Dinaledi Chamber to date is 1.55 $m^2$ and in the Hill Antechamber is 0.75 $m^2$. *Homo naledi* skeletal and dental material has come from these excavated areas and from collection of skeletal remains from the floor surfaces of these chambers. That material now numbers more than 2500 pieces, many of them small fragments but including a large number of complete and well-preserved elements. Nonetheless much of this skeletal material is highly fragile with loss of elements with thin cortical bone or substantially trabecular bone content. Our excavations in 2013 and 2014 in the Dinaledi Chamber identified buried skeletal material with articulated limb, cranial, and vertebral elements (Dirks et al 2015; Berger et al 2015) including portions that could be attributed based on spatial and developmental data to a partial juvenile skeleton (Bolter et al 2020). This extensive evidence of skeletal morphology and spatial positioning of material has informed our more recent work in the system in a variety of ways (Elliott et al, 2021).

In many paleoanthropological contexts, the recovery of morphological or developmental observations on hominin skeletal material is a very high research priority. Additionally, some settings pose taphonomic questions that can be addressed only through collection of data from bone surfaces. To recover such data, hominin skeletal material has often been subjected to intensive preparation, cleaning, and consolidation. Much *Homo naledi* skeletal material from our earlier work has been treated with conventional methods of skeletal conservation and analysis, which has included microscopic examination of bone surfaces (Dirks *et al.,* 2015). Having this published evidence in hand gives us the possibility of taking a less destructive approach to studying the burial features that we identified in subsequent work. For these reasons, once we identified the possibility that the Dinaledi Feature 1 and Hill Antechamber Feature were burial contexts, we planned for non-destructive methods to record and preserve the spatial relationships between any remains entombed within these features. In the Hill Antechamber case, this involved lifting the feature *en bloc* for study, scanning, and possible preparation in the laboratory.

In the case of Dinaledi Feature 1, the prior recovery of a similar burial feature from the Hill Antechamber argued for leaving this feature in situ within the cave system and recording data on its spatial and sedimentary context noninvasively.

**Supplementary Information 4.1: Identification and assessment of skeletal remains from Dinaledi Feature 1**

Here we list brief anatomical identification of all catalogued skeletal elements and fragments excavated within and above the area of Feature 1. These specimens all come from grid units S950W550 and S900W550. Some other material excavated from unit S900W550 is associated with Feature 2 and these are listed separately below.

These brief identifications do not focus on comparative morphology beyond the identification to element where possible. Many of the remains are small fragments that are not identifiable to element and these are included here to present a complete record. In the few cases where diagnostic morphology of *H. naledi* is present, this is indicated. Where evidence for developmental stage is present, this is indicated.

We have not fully cleaned sediment from this skeletal material or applied fixatives or other chemicals, in anticipation of possible analysis of biochemicals or trace evidence. This precludes a full examination of possible surface modifications or other surface taphonomy.

Nearly all elements identified in this list are compatible with belonging to a single adult individual. All identifiable adult elements are size-compatible; they do not appear to represent a mixture of individuals of different sizes. None of the adult elements duplicate each other; nor is there any duplication with identifiable elements that remain *in situ* within Feature 1 in the Dinaledi Chamber.

There are three notable exceptions that are immature elements and inconsistent with the single adult represented by the rest of this material. Two of these are fragments of juvenile femur, including the right proximal femur fragment included in U.W. 101-2250 and a fragment of femur shaft U.W. 101-2260. These two elements were excavated overlying and in physical contact with the bone concentration of Feature 1, both within 5 cm of each other near the center of the feature. The proximal right immature femur duplicates the proximal right adult femur fragment, also included within U.W. 101-2250. These two elements were excavated in immediate contact with each other with the immature femur fragment overlying the adult fragment. The other inconsistent element is an immature proximal left humerus fragment, U.W. 101-2243. This element duplicates another proximal left humerus within Feature 1, U.W. 101-2231. These two elements contrast in developmental status, with U.W. 101-2243 having evidence of an unfused proximal epiphysis and U.W. 101-2231 compatible with adult status. The two elements are also in different situations relative to the feature: U.W. 101-2231 is near other compatible humerus and upper limb material at the south end of the feature, while U.W. 101-2243 was out of anatomical placement overlying the uppermost part of the north side of the feature. We interpret these three fragments of immature elements as bone fragments that were on the surface or within pre-existing deposit that comprised the sediment fill of Feature 1.

**Specimen list**

U.W. 101-2114

Thin plate-like bone fragment, broken on all edges, less than 15 mm.

U.W. 101-2115

Two cranial bone fragments. One is possibly zygomatic or root of zygomatic arch on temporal, length 23.8. The other is possibly zygomatic at orbit, or frontal above orbit, bone surface is concave and slightly wavy. Preserved size 16.5 x 16.5.

U.W. 101-2116

Phalanx fragment with adhering clump of sediment. Fragment size is 18.9 x 9.0

U.W. 101-2117

Right proximal radius fragment. The head is present but broken around the visible edge, and one side is still obscured by sediment. The bone is very comparable in size with U.W. 101-2246 and they may be antimeres. 38.8 mm length

U.W. 101-2118

Two bone fragments less than 15 mm

U.W. 101-2119

Bone fragment less than 15 mm

U.W. 101-2120

Two cranial bone fragments less than 20 mm

U.W. 101-2121

Right proximal radius fragment, representing nearly the entire circumference of the bone. The head is not present. The distal break on this fragment is consistent with the break at one end of U.W. 101-2240 and may be the same bone. Preserved length 49.6. Shaft diameter 10.0 x 9.4.

U.W. 101-2136

Cranial or mandibular fragment, possibly zygomatic arch. Length 20.4

U.W. 101-2137

Three large clumps of sediment with spongy bone fragments embedded within them. Possibly distal femur or pelvic.

U.W. 101-2138

Sediment clump with embedded bone fragments. ca. 15 mm

U.W. 101-2139

Sediment clump with embedded fragments of cortical bone. ca. 15 mm

U.W. 101-2140

Tooth root. 11.5 mm

U.W. 101-2141

Tooth root. 13.2 mm.

U.W. 101-2142

Cortical bone fragment, 16.0 mm.

U.W. 101-2143

Three bone fragments less than 15 mm.

U.W. 101-2144

Mandibular incisor. Probably left I1, possibly I2. Occlusal wear is extensive and remaining crown height above cervix on distal face is only 2.2 mm. The wear is at an angle of approximately 60 degrees compared to labial face. The wear has exposed dentin with MD 2.0 mm, LL 1.3 mm. LL breadth 5.9, MD 4.6. Root is nearly complete with small section of distal tip missing due to fracture.

U.W. 101-2145

Bone or dentin fragments, less than 5 mm, with some sediment.

U.W. 101-2146

Bone fragments in sediment clump, less than 10 mm.

U.W. 101-2147

Bone fragment with large cancellous structure in sediment clump. Less than 20 mm.

U.W 101-2148

Two bone fragments with sediment adhering. Smaller fragment 13.0, larger 21.3.

U.W. 101-2168

Sediment with bone fragments less than 5 mm.

U.W. 101-2170

Tibia shaft portion, 121 mm in length, represents midshaft. Consistent with adult. Diameter at point where posterior shaft flattens is 21.4 x 17.2. Other small shaft fragments, all less than 20 mm.

U.W. 101-2171

Long bone shaft fragment, 34.3 x 10.4

U.W. 101-2172

Fourteen long bone shaft fragments. Most represent a part of the bone circumference and are between 30 and 40 mm in length, 10-15 in width. Additional fragments in bag.

U.W. 101-2173

Two bone fragments less than 15 mm.

U.W. 101-2174

Mandibular fragments. Two tooth crowns are present with their roots, and three additional root fragments. The two crowns are both left mandibular molars. One is markedly more worn, with large dentin pools at protoconid and endoconid, and metaconid is entirely fragmented away. The rest of the enamel rim has large fragments missing. BL 11.1 MD 12.1. This tooth has a distal interproximal facet, I interpret it as an M2. The other tooth is worn approximately flat with dentin exposed on mesial four cusps. BL 10.8 MD 12.1. This tooth is more triangular in shape distally and does not have an interproximal facet distally; I interpret it as an M3. One of the broken root fragments is a molar root, the others appear likely to be premolar roots. The morphology and dimensions of these teeth are compatible with *H. naledi*.

U.W. 101-2175

Lower left fourth premolar. Root is largely intact, broken at tip. Single root. Enamel is worn flat with large dentin pools for both buccal and lingual cusps. BL 10.7, MD 9.2. The morphology and size of this tooth are compatible with *H. naledi*.

U.W. 101-2176

Lower left third premolar. Two roots, both intact. BL 10.6, MD 9.4. Crown is worn flat, dentin exposed at protoconid and metaconid. This tooth possesses diagnostic anatomy for *H. naledi*.

U.W. 101-2177

Ulna shaft portion, 48 mm.

U.W. 101-2178

Two fragments. One is fibula shaft, 37.5 mm. The other is a flat fragment less than 20 mm.

U.W. 101-2179.

Bone fragment, possibly tarsal, maybe medial cuneiform fragment. Two additional pieces are bone fragments less than 10 mm.

U.W. 101-2180

Distal ulna shaft fragment, 44.6 mm.

U.W. 101-2181

Cranial fragment consistent with right zygomatic bone including orbital border. 34.8 x 24.7 mm.

U.W. 101-2182

Fibula or metacarpal shaft portion, 27 mm

U.W. 101-2183

Radius or humerus shaft portion 43.6 mm. Additional fragments of shaft.

U.W. 101-2184

Base of right mandibular corpus. 52.5 mm.

U.W. 101-2185

Enamel fragment, 5 mm.

U.W. 101-2186

Enamel fragment, corresponding to rim of molar, 8.9 mm

U.W. 101-2187

Tooth root fragment, 10.1 mm

U.W. 101-2188

Cranial vault fragment. One side is obscured by sediment. 21.4 x 15.4 mm, 5.4 thick.

U.W. 101-2189

Mesial part of mandibular molar, with mesial root complete. Very little enamel on this crown. This looks like a mirror image of the U.W. 101-2174 M3 mesial root. Several additional cranial or mandibular fragments, mostly less than 20 mm.

U.W. 101-2190

Fragment of humerus head, with small slice of surface 22.0 x 8.3 mm.

U.W. 101-2191

Partial phalanx, probably manual intermediate phalanx. Proximal articular surface is eroded but present, shaft 16.5. Embedded in sediment chunk.

U.W. 101-2199

Phalanx shaft portion, or possibly fibula shaft portion. 23.5 mm.

U.W. 101-2200

Fibula shaft or metatarsal shaft portion, triangular cross section. 22.6 mm

U.W. 101-2201

Three bone fragments, with some articular surfaces.

U.W. 101-2202

Bone fragment, shaped like possibly zygomatic or mandibular fragment, seems too thick to be vertebral or rib. 22.7 mm length.

U.W. 101-2203

Shaft fragment consistent with metacarpal. Length 28.5, diameter 5.9 x 5.2.

U.W. 101-2204

Bone fragment. 25.8 long, 14.8 broad, 6.8 thick.

U.W. 101-2205

Long bone shaft fragment, based on size and curvature consistent with humerus, representing approximately 40% of the circumference. Length 43.5, width 13.3.

U.W. 101-2206

Bone fragment, one side exposed trabeculae, the other rough cortical surface. Possibly distal femur or ilium. 24.5 x 17.9

U.W. 101-2207

Metatarsal or manual phalanx shaft fragment. 23.6 long, 7.9 x 5.3 near base.

U.W. 101-2208

Long bone shaft fragment, based on size and curvature probably humerus, representing 30% of the circumference. 25.4 x 13.9.

U.W. 101-2209

Long bone shaft fragment, based on size and curvature probably humerus, representing 30% of the circumference. 29.1 x 11.4.

U.W. 101-2210

Bone fragments, probably cranial, less than 30 mm.

U.W. 101-2211

Rib fragment. 34.3 long, diameter 8.6 x 6.1

U.W. 101-2221

Bone fragments.

U.W. 101-2222

Cranial vault fragment. 22.4 x 14.8.

U.W. 101-2223

Long bone fragment. 38.7 x 13.0

U.W. 101-2224

Left distal humerus fragment. The lateral supracondylar crest is well preserved and evident. Length 54.4.

U.W. 101-2225

Long bone shaft fragment with rounded circumference compatible with humerus or radius, including 50% of shaft circumference. 29.9 long, 12.0 wide.

U.W. 101-2226

Flat long bone shaft fragment. Consistent with distal humerus or proximal tibia. 38.5 x 15.8.

U.W. 101-2227

Long bone shaft fragment consistent with humerus. 29.1 x 15.1

U.W. 101-2228

Shaft fragment consistent with small long bone, metacarpal or metatarsal. 21.3 x 9.5

U.W. 101-2229

Long bone shaft fragment consistent with radius, ulna, or fibula. 27.4 x 10.2

U.W. 101-2230

Bone fragments in sediment.

U.W. 101-2231

Proximal left humerus shaft. Length 53. Diameter at surgical neck 18.2 x 15.8. A chip of enamel is adhering within the sediment that fills the proximal end of this fragment. The enamel chip is approximately 3 mm x 5 mm and could be molar or incisor. A second fragment refits the distal end of the first. Length 24.8 x 13.7.

U.W. 101-2232

Bone fragment 19 x 9 mm.

U.W. 101-2233

Bone fragment 18.7 x 12.8.

U.W. 101-2234

Bone fragment 20.6 x 8.4.

U.W. 101-2235

Tooth root, slightly bilobate toward crown. 15.1 mm.

U.W. 101-2236

Bone fragment with concave surface exposed on one side, other side trabecular fragments with sediment obscuring detail. 21 x 17.

U.W. 101-2237

Maxillary left molar. BL 12.3, MD 11.6. All roots are present and intact, all three curve distally, and the buccal two roots curve into each other strongly distally. Based on roots and crown morphology this resembles an M3 more than either other molar. The crown is bigger than any of the M1s and those teeth do not have the posteriorly directed lingual root that this one has. Occlusal surface has slight wear, no dentin exposure and all cusps are salient.

U.W. 101-2238

Mandibular ramus fragment of right coronoid process. 35.9 x 14.8.

U.W. 101-2239

Shaft fragment of femur or tibia, relatively flat, consistent in size and thickness with U.W. 101-2259 femur. Length 48.3, width 19.4.

U.W. 101-2240

Shaft portion of radius or ulna. 43.1 long, diameter obscured by sediment. This fragment is consistent with the break at the distal end of the U.W. 101-2121 radius fragment and may be the same bone. A small thin and slightly rounded bone fragment was adhering to one end, not in anatomical position. This is thinner than the shaft fragment and probably belongs to some other bone.

U.W. 101-2241

Bone fragment, thin with adhering sediment on one side. 19.9 x 13.6

U.W. 101-2242

Proximal right ulna, lacking olecranon process. Length 50.2, diameter below coronoid process 12.9 x 12.4. Two additional shaft portions refit the larger fragment.

U.W. 101-2243

Proximal left humerus shaft. The proximal end of this includes a small portion of metaphyseal surface across the lateral 25% of this end of the bone. There is additionally a small (10 mm) piece of epiphysis that was adhering in position to part of this lateral edge of the proximal end. This is now detached and its surface does correspond to the opposing surface of the diaphysis. Length 46.7. Shaft diameters at surgical neck 18.5 x 15.5.

U.W. 101-2244

Proximal end of a rib with the articular part of the head missing but the tubercle present. The neck is round in cross section and the break just lateral to the tubercle has a rib-like cross section. Length 25.9, diameter 6.3.

U.W. 101-2245

Bone fragment consistent with vertebral lamina. 13.3 x 9.3 x 5.6 thick at thickest point.

U.W. 101-2246

Proximal left radius, lacking head. Radial tuberosity and approximately 25% of shaft are present. The neck is broken and does not preserve a metaphyseal surface. It is size-consistent with adult material. That makes it comparable to the humerus elements nearby including U.W. 101-2243. Length 55.7, diameter of neck 10.1 x 9.2.

U.W. 101-2247

Phalanx or rib fragment. 20.6 x 8.2 x 4.9.

U.W. 101-2248

Rib fragments less than 20 mm.

U.W. 101-2249

Right distal humerus, lacking trochlea and capitulum. Length 88. Diameter above epicondylar ridges 14.1 x 15.7.

U.W. 101-2250

Long bone remains that represent at least two different elements. One of these is a right proximal femur, including base of neck, lesser trochanter, broken below greater trochanter. Subtrochanteric diameters AP 20.9, ML 28.5. Length of fragment 68.4. A second large fragment is a portion of long bone shaft, 62.5 long, 16.0 in diameter. The cortical bone is thin relative to the shaft diameter, suggesting that it is more consistent with juvenile femur or tibia rather than adult humerus. In size, this shaft fragment may be compatible with the immature proximal femur fragment U.W. 101-2260 but there is no refit between these pieces. This bone is highly dark-stained with iridescent sheen and edge. Smaller additional fragment of same 27.3 x 14.0. Additional bag contains bone fragments in sediment.

U.W. 101-2258

Rib fragment 23.1 long.

U.W. 101-2259

Femur shaft. Refits right proximal femur in U.W. 101-2250. Fracture is stained, not fresh. Length 152.9. Diameter same as 101-2250.

U.W. 101-2260

Immature proximal right femur. Includes neck, head is missing. It is possible that the metaphyseal surface for head is present here but sediment and possible erosion mask whether this is the case. The metaphyseal surface for the greater trochanter is partially present. None of the head is present, and the broken portion of the neck does not retain any of its metaphyseal surface. Lesser trochanter is projecting, shaft is broken away irregularly and lateral border of shaft missing. Fragment 41 mm, neck 19.6 x 15.1.

U.W. 101-2261

Fragment with concave surface on one side, convex obscured by sediment on the other. Fragment 19.7 x 15.2 x 6.6 thick.

U.W. 101-2262

Long bone shaft fragment, ulna. 39 x 13.4.

U.W. 101-2263

First metacarpal head and 40% of shaft. Includes diagnostic morphology of *H. naledi.* 28.5 mm.

U.W. 101-2264

Shaft fragment of phalanx or metacarpal. 25 mm x 8.1 mm.

U.W. 101-2265

Bone fragment, possibly carpal fragment less than 15 mm.

U.W. 101-2266

Bone fragment, less than 20 mm.

U.W. 101-2267

Bone fragments within clump of sediment, possibly rib fragments, 32.7 mm.

U.W. 101-2268

Bone fragment, possibly rib fragment, 25.4 x 9.4.

U.W. 101-2269

Bone fragment with morphology and possible metaphyseal surfaces, possibly immature long bone fragment? 18.4 mm.

U.W. 101-2270

Bone fragments

U.W. 101-2271

Bone fragment less than 20 mm.

U.W. 101-2272

Long bone fragment 25.2 x 16.6 x 6.9 thick.

U.W. 101-2273

Shaft fragment of long bone, 28.5 x 17.4. Other associated fragments.

U.W. 101-2274

Manual phalanx. Head is eroded, base is missing. Probably proximal phalanx based on length. 28.0 mm long, diameters at midshaft 8.8 x 4.9.

U.W. 101-2275

Ischium fragment including lunate surface of acetabulum, broken superior to ischial tuberosity. 25.0 x 13.0 mm of lunate surface present. Fragment length 21.1 from acetabular border to inferior edge. Subacetabular sulcus 6.9 mm.

U.W. 101-2276

Mandibular corpus fragment, including the angulation between the base of the corpus and either the external or internal surfaces. Based on curvature this seems likely to be external. This fragment is possibly consistent with U.W. 101-2184, which is clearly right mandibular corpus including the base and the swelling at the base of the ramus. Fragment dimensions 36.5 x 13.5.

U.W. 101-2277

Shaft portion of ulna or fibula. Length 37.3, diameter 9.9 x 9.4.

U.W. 101-2278

Left proximal ulna with shaft fragments. Olecranon process is present but eroded, coronoid process appears to be broken or missing. The coronoid process is certainly missing because of a break. This is consistent with adult ulna due to the extent of the olecranon process that remains without evidence of metaphysis, but its most proximal extent is abraded. Measurements below coronoid 11.9 x 11.5. Length 47.6.

#### Supplementary Information 4.2. Identification and assessment of skeletal remains from Dinaledi Feature 2

Very little skeletal material has been excavated from or above Dinaledi Feature 2. The feature remains largely undisturbed, with only a small semi-circular concentration of bone visible and an unknown portion remaining within the unexcavated S950W600 grid square. The identifiable elements are consistent with a single adult individual but very little evidence has been recovered to date.

U.W. 101-2134

Bone fragment, thin cortical flake, 13.9 mm

U.W. 101-2135

Bone fragment consistent with cranium or mandible, 12.7 mm

U.W. 101-2166

Bone fragment consistent with cranium or mandible, 34.9 x 20.9 mm.

U.W. 101-2167

Bone fragment in sediment, possibly phalanx shaft. Less than 20 mm.

U.W. 101-2198

Flat thin fragment consistent with zygomatic arch or vertebral lamina. 15.2 x 7.4.

U.W. 101-2220

Femur or tibia shaft portion. Surface morphology and cross-section obscured by sediment.

Length 80.7, diameter 24.9

### Supplementary Information 5: Postmortem change and *Homo naledi*: Understanding decomposition in the Rising Star burial environment.

Decomposition can be described as the process by which the body physically breaks down and decays, ultimately resulting in skeletonization. However, it is important to firstly understand the nature of body as it decomposes, as although the distal outcomes (skeletonization and post-skeletonization phases) may be different in differing depositional scenarios, the initial proximal stages (the early postmortem period) are entirely driven by the linked processes of putrefaction and decomposition. Decompositional changes to a body that occur immediately after death are more rapid than those occurring later in the process, making it difficult to establish an exact time interval, but the general pattern of decomposition is well understood (Bristow et al., 2011; Gill-King, 1997; Wilson et al., 2007). The factors that affect the rate at which postmortem changes occur are classified into two groups (Prieto et al., 2004); firstly, those dependent on the cadaver and intrinsic physiological factors, and secondly, those dependent on the postmortem environment of the cadaver and extrinsic factors such as temperature, humidity, insect activity, and scavenging with the codicil that decomposition can be retarded by several processes, including physical and chemical barriers and climatic factors (Bristow et al. 2011). Much of our understanding of human decomposition data is derived from forensic taphonomic research either based on real-world forensic case evidence and interpretation, or (more crucially) actualistic experimentation undertaken at human taphonomic facilities - such forensic approaches override classical inductive and intuitive palaeo and archaeo-taphonomic approaches which come with significant evidential shortcomings (Bristow et al. 2011; Schotsmans et al., 2022). Whilst the general process of decomposition is broadly similar across most mammalian taxa, specific differences in functional anatomy, body composition, physiology, biochemistry, microbiome, and skeletal ultrastructure (including histological pattern, mineralized and organic makeup, and biomechanical competency) between the human pattern (and by inference hominin and non-human primate) and that of other animals, limits the use of inference from classical vertebrate taphonomic studies of animal decomposition and skeletonization (Bristow et al 2011).

Decomposition processes are influenced primarily by temperature, but secondarily affected by many different variables, including humidity, insect activity, sunlight, rainfall, scavenging, and the burial (sedimentary) environment, among others (Bristow et al. 2011; Henderson, 1987). Temperature influences microbial degradation as the primary driver of decomposition (Forbes et al., 2017; Ody, et al., 2017) with warmer temperatures increasing the rate of mircrobially-driven decomposition. In bodies exposed on the external ground surface (sub-aerially) insect activity is the key driver of soft tissue decomposition in conjunction with microbial action. Very cold or freezing temperatures inhibit and retard microbial action leading to tissue preservation or the drastic slowing of the decomposition process. Low temperatures and/or enclosed spaces (including deep cave systems such as Rising Star) also influence insect action and access - holometabolous insects (those with larval and pupal stages) are either inactive or have limited activity (Anderson & Cervenka 2002; Bachmann and Simmons 2010; Benecke et al 2008; Komar, 1998; Lutz et al. 2021; Michaud and Moreau 2009; Mona et al 2019; Myskowiak, et al 1999.).

It is important to understand the physical changes that the body (cadaver) undergoes between death and skeletonization, as these changes influence both the volume and integrity of body systems and parts (Boulestin and Duday 2005; Duday, 2009; Schotsmans, et al., 2022). The process of decomposition proceeds through six recognized stages (after Wilson et al., 2007): 1) Fresh - begins at death, and includes rigor mortis, postmortem hypostasis and cooling which continues until bloating of the cadaver is visible; 2) Primary bloat stage – bacterially mediated accumulation of gases within the body causes distension and bloating, with an increase in surface area and volume of the body. Whilst there is no disarticulation of body parts, the hair and epidermis are loose, and the soil-skin interface is visually and biochemically altered by microbial-sediment interaction. The body emits a strong odour caused by release of volatile organic compounds (primarily putrescene and cadaverine); 3) Secondary bloat stage – the body is still bloated, and there is now rank-order disarticulation of joints and limb segments. Purging of decomposition fluids occurs from natural cavities such as the oral and nasal cavities, anus, or from damage to the integrity of the integument. The soil-skin interface is black. A mobile leachate plume may extend into the surrounding soil/sediment around and some distance from the body; 4) Active decay stage - deflation of the cadaver, disarticulation of joints, limbs and head. Muscle and integument may still be present. The cadaver is very wet with a strong odour; 5) Advanced decay stage - collapse of components of the rib cage and pectoral gridle into the thoracic cavity, with most of the flesh liquified or gone; skin, bone, fat, and cartilage may remain and the cadaver and underlying sediments are very wet with leachates and decomposition fluids; and, 6) Skeletonization – muscles, skin, fat and cartilage disappear, though some persistent ligaments may remain. Bone will present as ‘dry’ with a shift in biomechanical competency through the loss of organic (primarily collagen) content, and chemi-absorbed and physi-absorbed water, leaving the mineral bioapatite as the primary component of remaining skeletal tissue.

Bone will facture in a classic ‘postmortem’ and opposed to ‘perimortem’ fashion (Lyman, 1984; Christensen et al 2022).

In the early post-mortem period, enzymatic digestion (autolysis) is the primary driving factor. This initiates widespread cell degradation via anaerobic microorganisms such as bacteria, fungi, and protozoa (Vass, 2001, Vass, 2011, Vass, et al., 2002) within the gastrointestinal tract and the respiratory system (Carter, et al., 2007, Carter, et al., 2010, Carter, et al., 2008a&b), leading to saturation of body tissues (haemolysis) (Fielder and Graw, 2003) with decomposition products. These products result in colour changes seen on the dermis with concomitant bloating of the body, signaling the onset of the putrefactive stage of decomposition (Bristow, et al., 2011). A continued internal build-up of decompositional gases eventually leads a build-up of internal pressure, leading to purging through body orifices, such as the mouth, nose, and anus, and possible tissue rupturing (Carter, et al., 2007). The active decay process is most often associated with a decrease in body mass (Adlam and Simmons, 2007). Putrefaction processes function under anaerobic conditions (Adlam and Simmons, 2007, Cross and Simmons, 2010, Simmons, et al., 2010a, Simmons, et al., 2010b). Further decay by bacteria and fungi is purely anaerobic and it is this which leads to full skeletonization of the cadaver (Fielder and Graw, 2003). The decay process will generally be accelerated in bodies in sub-aerial contexts, particularly with open access to insect faunas. Different depositional and burial environments will lead to a difference in observed or expected results since neither environment is static (Wilson, et al., 2007).

In burial environments, there is a restriction of access to the remains from scavengers and insects who pose a major role in decomposition (Adlam and Simmons, 2007, Cross and Simmons, 2010, Simmons, et al., 2010a, Simmons, et al., 2010b), as well as restricting the effects of sub-aerial processes such as weathering, which can accelerate the decomposition and degradation of skeletal tissue itself (Beherensmeyer, 1978, Hill, 1976; Manhein, 1996; Tappen, 1969). Most of the scientific literature regarding the direct observations of decompositional processes of human remains refers to those that have been deposited on the surface of the ground (refs), or more recently in shallow or open pits (e.g. Mickleburgh and Wescott, 2018; Mickleburgh et al., 2022), where temperature and insect activity play significant roles in the decomposition process (e.g. Megyesi et al., 2005). In burial environments where the body is encapsulated, such as inside structures or buried in sediment matrix, there is restriction of access to the remains from scavengers and insects who would otherwise play a major role in the pattern and tempo of decomposition (e.g. Cross, et al., 2010; Simmons et al., 2010a&b). In sub-surface decomposition soil temperature (Carter and Tibbett 2006; Carter et al. 2008a), moisture content, soil texture and type (Fielder and Graw 2003; Tibbett et al. 2004), soil pH (Haslam and Tibbett 2009; Prangnell and McGowan 2009) and bacterial/microbial community structure (Carter and Tibbett, 2006 and 2008; Hopkins, 2008; Hopkins et al., 2000; Sagara et al., 2008; Zhang et al., 2021) all play an important role in the postmortem fate of the buried cadaver.

During and following the process of skeletonization the body undergoes a process of disarticulation and scattering of elements, due to the decomposition of muscles, connective tissues (particularly ligamentous structures, but also tendons) and joint capsules. The process of disarticulation specifically relates to the destruction, decomposition, or removal of (primarily) soft tissues which hold bony elements or joint surfaces of a skeleton together. Depending on the organism under consideration (and anatomical region) these soft tissues may comprise tendons, ligaments, muscles or skin, or components of joint capsules such as synovial membrane; the breakdown of these anatomical components allows individual bones, or complete elements and limbs, to disarticulate from the body. Because soft tissue anatomy varies with the functional anatomy of each joint, the process of disarticulation is highly complex (Roksandic, 2002). Subsequent dispersal is the increase or the decrease of the distance between bones (Duday and Guillon 2006). The movement of bones out of normal anatomical association (termed necrodynamics; Mickleburgh and Wescott, 2018) occurs through the combined factors of connective tissue and joint decomposition, the effects of gravity on the bones, physical disturbance, and sediment type, grain size and stability. Movement can be affected by both physical processes (including swelling and contraction of sediment under moisture cycles; after Pokines et al., 2018), biotic agents (animal and plant bioturbation; Armour-Chelu and Andrews, 1994; Gabet et al., 2003; Pokines and Baker, 2013) or human agents (Hunter and Cox 2005). Scattering can be extensive and spatially widespread in open sub-aerial contexts due to the transport effects of gravity and water (Haglund, 1993), or scavengers (Berryman, 2002; Carson, et al.., 2000; Haglund et al., 1989; Haynes, 1982) or minimal, where skeletal elements and body units (head, thorax, appendicular skeleton etc) are supported and encapsulate by matrix such as soil, volcanic ash, or anthropogenic materials such as concrete (after Hunter and Cox, 2005; Pokines and Baker, 2013). Displacement of body elements requires the presence of open space for the bones to move within and into. Such spaces are termed primary where the empty space is the burial chamber itself (which can include mortuary structures, coffins, pits, or surfaces), and secondary spaces generated by the decomposition of soft tissues (Duday, 2009).

However, the order and pattern of rank disarticulation in the human (hominin) body differs somewhat from that observed in quadrupedal animals (after Lyman, 1994), but has been difficult to quantify precisely. Initial observations based on an understanding of musculo-skeletal anatomy and biomechanical joint function led researchers (Duday, 2005 and 2009; Knusel, 2014; Knusel and Robb, 2016) to define two primary types of articulations termed persistent (durable) and labile (non-durable). Persistent joints are considered to be those which are mechanically stable and strong, and which play a role in important biomechanical functions such as weight bearing and locomotion (such as the atlanto-occipital, humeroulnar, sacro-iliac, and tibio-tarsal joints, and the structures of the thoracic and lumbar vertebrae). Labile joints are considered to be those prone to rapid decomposition due to a lack of stabilizing and regulating bony articulations or soft tissue support (such as those between cervical vertebrae, carpal-metacarpal-phalangeal joints, the costosternal and costovertebral joints, scapulothoracic joint, and tarsal-metatarsal-phalangeal joints). Subsequent studies (Duday, 2009; Knüsel, 2014; Knüsel and Robb, 2016) have indicated that some joints were misclassified (i.e. femoroacetabular) and that the relationship between biomechanical competency in life and the persistence of such joint structures after death is not straightforward or simple. Current consensus suggests that persistent joints are represented by the atlanto-occipital, humeroulnar, thoracic and lumbar inter-vertebral, lumbosacral, sacroiliac, tibiofemoral, talocrural, and talocalcaneal joints. Labial joints are represented by the hyoid and its anchoring attachments, temporomandibular, cervical vertebral, scapulothoracic, glenohumeral, Dcostosternal, costovertebral, acetabulofemoral, femoro-patella, carpal, metacarpal, tarsal, metatarsal, and phalangeal joints (though see Schotsmans et al., 2022: 512 for a summary of ongoing disagreements in classification).

In the practice of archaeothanatology (Duday, 2005 and 2009; Knüsel, 2014; Knüsel and Robb, 2016) it is the differentiation between articulations of labile and persistent joints that is used to distinguish primary and secondary burials. Archaeothanatology uses the relative sequence of joint disarticulation to separate natural (biotic and abiotic) processes from those relating to the placement and treatment of the body (such as rapid primary burial). However, both archaeothanatological and forensic actualistic studies have indicated that patterns of joint disarticulation and final bone position covary with depending on original body disposition/placement as well as secondary environmental effects (Haglund, 1993; Rodriguez and Bass, 1985; Gerdau-Radonic, 2012). The overall pattern suggests that disarticulation proceeds in a generally craniocaudal direction, and from the extremities to the core (from the distal appendages to the thorax). Onto this must be mapped the specific patterns of association, disarticulation, rotation and displacement of each joint, if an understanding of the postmortem narrative is to be achieved. However, based on actualistic experimental forensic taphonomy Schotsmans and colleagues (2022) suggest that labile and persistent joints disarticulate over different time intervals, irrespective of their classification, and that joint disarticulation is complex and influenced by many covariables, with the pattern of disarticulation being highly influenced by slight differences in body position.

**Supplementary Information 5.1: Spatial taphonomy of Puzzle Box – archaeothanatology of Hand and Foot 1.**

In situ sequential scanning of the fossil deposits in the Puzzle Box was undertaken during the initial excavations of the Dinaledi Chamber (2013 and 2014). This was accomplished with structured light scanning to assist in spatial recording and visualisation of the position and disposition of individual bones. The technical methods and workflow employed are detailed Kruger and colleagues (2015). In particular the recording methodology provides significant spatial taphonomic information about the decomposition of body parts within the Puzzle Box burial environment. Most informative are Dinaledi Hand 1 and Foot 1. Dinaledi Hand 1 (H1) is a nearly complete right hand, found semi-articulated with the palmar surface facing upwards with the bones in close spatial association. Dinaledi Foot 1 (F1) is a semi-articulated adult right foot, found resting on its dorsal (plantar) surface.

Scanning of the fossils within their sediment matrix was enacted with a handheld Artec Eva 3D surface scanner. A series of sequential scans were taken of H1 and F1 prior to excavation, and then at different stages during the excavation process as sediment was removed, prior to recovery of the fossils. Following excavation, lifting, cleaning and conservation, each bone was surface scanned using a NextEngine 3D Laser Scanner, with data exported as ply format files. The individual bone scans were then imported into Artec Studio 10 Professional along with the sequential surface scans undertaken during excavation. Translation and rotation of each 3D model of the individual hand bones was guided and aligned by surface models of the excavation. By layering each sequential scan of the in-situ hand and foot, a virtual three-dimensional representation of the excavation area surrounding H1 and F1 was produced, representing a cube of sediment within which the precise anatomical resting and relative position of each bone element can be seen. The encapsulating sediment was then extracted from the scan volume, leaving the precise spatial arrangement of each element as they would have been in the ground, prior to excavation and recovery. This 'reverse engineering' of the fossil deposits allows for the investigation of spatial taphonomic patterns within the surviving assemblage, as well as clear visualisation of small-scale anatomical articulations and disjunctions between bones. This produced an accurate 3D reconstruction of the placement of H1 and F1 as they lay in-situ, by virtually placing the bones of H1 and F1 back into the excavation pit. This allows for a precise and accurate visualisation of the relative position of each elemental part, which imparts significant spatial information which can assist in taphonomic analyses and interpretation (supplementary figure S20).

The extracted (sediment-removed) anatomical volumes indicate that both Hand 1 and Foot 1 retain their labile joint structures. In particular, H1 was almost completely articulated, and the reconstructed hand indicates that it was flexed (closed) or semi-closed during the process of skeletonization. The overall configuration of F1 suggests that it was resting on the dorsal surface. To retain such fragile labile anatomical structures observed in H1 and F1 is most parsimoniously explained by rapid encapsulation within the burial environment, with the position of individual structures supported by sedimentary matrix as the soft tissues of the hand and foot decomposed – this matrix support would ensure little or no movement of the individual elements of the hand and foot during decomposition, supporting the interpretation of rapid burial.

#### Supplementary Information: Tables

Supplementary Table 1: Particle-size distribution (PSD) of sediments based on the Folk and Ward Method.

| Sample name | Folk and Ward Method (µm) | | | |
|---|---|---|---|---|
| | Mean grain size | Sorting | Skewness | Kurtosis |
| DF1 | 372.21 | 2.60 | -0.07 | 0.78 |
| DF2 | 279.24 | 3.22 | -0.07 | 0.85 |
| DF3 | 442.02 | 2.28 | -0.06 | 0.75 |
| DF4 | 373.84 | 2.90 | -0.23 | 0.91 |
| DF5 | 356.86 | 2.61 | -0.03 | 0.76 |
| DF6 | 444.55 | 2.28 | -0.08 | 0.75 |
| DF7 | 375.33 | 2.54 | -0.05 | 0.77 |
| DF8 | 433.52 | 2.29 | -0.05 | 0.74 |
| DF9 | 379.40 | 2.49 | -0.03 | 0.74 |
| DF10 | 443.43 | 2.27 | -0.06 | 0.74 |
| DF11 | 351.16 | 2.81 | -0.12 | 0.83 |
| DF12 | 336.17 | 2.68 | -0.02 | 0.74 |
| DF13 | 369.75 | 2.51 | -0.02 | 0.73 |
| DF14 | 416.46 | 2.34 | -0.04 | 0.73 |
| DF15 | 334.22 | 2.79 | -0.05 | 0.80 |
| DF16 | 306.38 | 2.89 | -0.02 | 0.77 |
| DF17 | 362.50 | 2.55 | -0.02 | 0.74 |
| DF18 | 369.84 | 2.52 | -0.02 | 0.73 |
| DF19 | 533.93 | 2.31 | -0.28 | 1.01 |
| DF20 | 385.75 | 2.46 | -0.03 | 0.74 |
| DF21 | 261.64 | 3.29 | -0.05 | 0.82 |
| DF22 | 305.33 | 2.87 | -0.01 | 0.75 |
| DF23 | 270.15 | 3.34 | -0.08 | 0.86 |

| | | | | |
|---|---|---|---|---|
| DF24 | 302.06 | 2.90 | -0.02 | 0.76 |
| DF25 | 340.90 | 2.82 | -0.08 | 0.84 |
| DF26 | 331.13 | 2.97 | -0.14 | 0.84 |
| DF27 | 208.47 | 3.89 | -0.07 | 0.84 |
| DF28 | 288.48 | 3.21 | -0.09 | 0.85 |
| DF29 | 317.93 | 2.87 | -0.05 | 0.80 |
| DF30 | 442.62 | 2.28 | -0.07 | 0.75 |
| DF31 | 268.14 | 3.32 | -0.07 | 0.85 |
| DF32 | 222.71 | 3.69 | -0.06 | 0.83 |
| DF33 | 337.60 | 2.71 | -0.03 | 0.76 |
| DF34 | 442.46 | 2.63 | -0.28 | 0.94 |
| SA1 | 323.81 | 2.84 | -0.05 | 0.80 |
| SA2 | 468.58 | 2.42 | -0.21 | 0.92 |
| SA3 | 277.86 | 3.27 | -0.09 | 0.85 |
| SA4 | 370.36 | 2.64 | -0.08 | 0.81 |
| SA5 | 345.26 | 2.66 | -0.03 | 0.76 |
| SA6 | 332.60 | 2.72 | -0.02 | 0.75 |
| SA7 | 308.77 | 2.84 | -0.01 | 0.75 |
| SA8 | 354.38 | 2.95 | -0.16 | 0.91 |
| SB1 | 237.81 | 3.51 | -0.05 | 0.82 |
| SB2 | 360.13 | 2.58 | -0.03 | 0.75 |
| SB3 | 358.12 | 2.67 | -0.06 | 0.80 |
| SC1 | 362.99 | 2.58 | -0.03 | 0.75 |
| SC2 | 328.85 | 2.81 | -0.05 | 0.80 |
| SC3 | 383.76 | 2.48 | -0.04 | 0.75 |
| SC4 | 332.60 | 2.70 | -0.02 | 0.75 |
| SE1 | 329.15 | 2.71 | -0.01 | 0.74 |
| SE2 | 265.82 | 3.36 | -0.09 | 0.84 |
| SE3 | 460.70 | 2.21 | -0.06 | 0.73 |
| SE4 | 385.72 | 2.48 | -0.04 | 0.75 |

| | | | | |
|---|---|---|---|---|
| SE5 | 278.21 | 3.25 | -0.08 | 0.84 |

Supplementary Table 2: Bulk major oxide chemistry and loss on ignition (LOI) obtained from x-ray fluorescence (XRF) in weight percentage (wt.%).

| **Sample name** | **Sample locality** | **Al2O3** | **CaO** | **Fe2O3** | **K2O** | **MgO** | **MnO** | **P2O5** | **SiO2** | **TiO2** | **LOI** | **SUM** |
|---|---|---|---|---|---|---|---|---|---|---|---|---|
| DF1 | DF group: Sediments from above Features 1 and 2 collected during excavation and opening of features. | 16.39 | 1.55 | 10.69 | 1.72 | 2.79 | 4.41 | 0.27 | 52.67 | 0.75 | 9.1 | 100.34 |
| DF2 | | 16.97 | 1.18 | 10.15 | 1.75 | 2.42 | 3.66 | 0.19 | 54.23 | 0.814 | 8.51 | 99.87 |
| DF3 | | 15.65 | 1.22 | 10.49 | 1.61 | 2.66 | 4.25 | 0.37 | 54.69 | 0.754 | 8.33 | 100.03 |
| DF4 | | 16.09 | 1.17 | 10.33 | 1.68 | 2.67 | 4.32 | 0.27 | 53.11 | 0.758 | 8.66 | 99.05 |
| DF5 | | 16.23 | 1.21 | 10.51 | 1.68 | 2.51 | 4.30 | 0.29 | 53.89 | 0.746 | 8.55 | 99.91 |
| DF6 | | 16.02 | 1.21 | 10.46 | 1.67 | 2.59 | 4.29 | 0.25 | 54.10 | 0.765 | 8.61 | 99.96 |
| DF7 | | 16.47 | 1.58 | 10.09 | 1.71 | 2.73 | 3.98 | 0.27 | 53.34 | 0.773 | 9.1 | 100.04 |
| DF8 | | 14.38 | 6.90 | 8.95 | 1.52 | 3.08 | 3.45 | 0.57 | 48.09 | 0.686 | 12.31 | 99.93 |
| DF9 | | 15.63 | 1.04 | 10.42 | 1.62 | 2.51 | 4.51 | 0.34 | 54.90 | 0.734 | 8.14 | 99.85 |
| DF10 | | 16.09 | 1.17 | 10.66 | 1.67 | 2.56 | 4.53 | 0.35 | 53.33 | 0.743 | 8.9 | 100.00 |
| DF11 | | 17.45 | 0.86 | 10.38 | 1.80 | 2.13 | 3.70 | 0.18 | 53.75 | 0.812 | 8.74 | 99.80 |
| DF12 | | 16.91 | 1.30 | 10.42 | 1.74 | 2.20 | 4.09 | 0.48 | 52.89 | 0.782 | 9.03 | 99.84 |
| DF13 | | 15.82 | 0.92 | 10.70 | 1.68 | 2.31 | 4.52 | 0.27 | 55.15 | 0.751 | 8.04 | 100.16 |
| DF14 | | 16.45 | 0.76 | 10.08 | 1.68 | 2.08 | 3.70 | 0.21 | 56.30 | 0.778 | 7.84 | 99.88 |
| DF15 | | 15.15 | 1.00 | 10.79 | 1.70 | 2.61 | 5.28 | 0.23 | 53.78 | 0.712 | 8.44 | 99.69 |
| DF16 | | 16.09 | 0.88 | 10.69 | 1.71 | 2.39 | 4.60 | 0.23 | 53.82 | 0.764 | 8.33 | 99.49 |
| DF17 | | 15.35 | 0.88 | 10.36 | 1.60 | 2.33 | 4.35 | 0.25 | 56.10 | 0.731 | 7.93 | 99.89 |
| DF18 | | 15.48 | 0.95 | 10.46 | 1.61 | 2.27 | 4.41 | 0.27 | 55.39 | 0.727 | 8.29 | 99.86 |
| DF19 | | 16.73 | 0.80 | 10.07 | 1.71 | 2.25 | 3.62 | 0.16 | 55.60 | 0.805 | 8.25 | 100.00 |
| DF20 | | 17.30 | 0.97 | 10.13 | 1.72 | 2.22 | 3.49 | 0.20 | 54.30 | 0.804 | 8.69 | 99.82 |
| DF21 | | 14.57 | 0.89 | 9.55 | 1.49 | 2.00 | 3.59 | 0.33 | 59.28 | 0.72 | 7.62 | 100.03 |
| DF22 | | 14.85 | 0.89 | 10.22 | 1.54 | 2.07 | 4.13 | 0.32 | 57.61 | 0.723 | 7.86 | 100.21 |

| | | | | | | | | | | | | |
|---|---|---|---|---|---|---|---|---|---|---|---|---|
| DF23 | | 14.13 | 0.96 | 9.78 | 1.42 | 2.32 | 3.86 | 0.34 | 58.71 | 0.702 | 7.65 | 99.86 |
| DF24 | | 14.69 | 1.00 | 9.50 | 1.42 | 2.33 | 3.45 | 0.39 | 58.44 | 0.719 | 7.6 | 99.53 |
| DF25 | | 15.02 | 0.91 | 9.51 | 1.44 | 2.36 | 3.50 | 0.29 | 58.57 | 0.743 | 7.67 | 100.01 |
| DF26 | | 14.78 | 1.08 | 9.28 | 1.43 | 2.32 | 3.20 | 0.40 | 58.76 | 0.757 | 7.61 | 99.62 |
| DF27 | | 15.18 | 0.98 | 9.54 | 1.52 | 2.12 | 3.74 | 0.36 | 57.51 | 0.763 | 7.73 | 99.43 |
| DF28 | | 13.71 | 1.01 | 9.31 | 1.35 | 2.18 | 3.78 | 0.31 | 60.38 | 0.702 | 7.27 | 99.99 |
| DF29 | | 14.80 | 1.55 | 9.96 | 1.52 | 2.22 | 4.60 | 0.75 | 55.51 | 0.701 | 8.05 | 99.66 |
| DF30 | | 14.26 | 1.18 | 9.64 | 1.47 | 2.02 | 4.02 | 0.51 | 58.44 | 0.707 | 7.5 | 99.74 |
| DF31 | | 16.19 | 0.88 | 10.10 | 1.69 | 1.95 | 3.88 | 0.30 | 56.21 | 0.781 | 7.96 | 99.94 |
| DF32 | | 14.88 | 1.36 | 9.80 | 1.53 | 1.94 | 3.90 | 0.63 | 57.03 | 0.734 | 7.7 | 99.51 |
| DF33 | | 14.10 | 2.11 | 9.45 | 1.49 | 1.91 | 3.92 | 1.16 | 57.55 | 0.692 | 7.49 | 99.87 |
| DF34 | | 16.09 | 2.03 | 10.19 | 1.68 | 2.01 | 3.54 | 1.14 | 54.58 | 0.784 | 8.05 | 100.11 |
| **Mean** | | **15.59** | **1.30** | **10.08** | **1.60** | **2.33** | **4.02** | **0.38** | **55.53** | **0.75** | **8.28** | |
| **STD** | | **0.98** | **1.04** | **0.48** | **0.12** | **0.28** | **0.45** | **0.23** | **2.50** | **0.04** | **0.87** | |
| SA1 | SA group: Sediment from sterile areas east of Feature 1 | 14.02 | 4.70 | 9.12 | 1.44 | 4.29 | 3.60 | 0.75 | 48.99 | 0.666 | 11.89 | 99.47 |
| SA2 | | 15.12 | 0.81 | 10.52 | 1.56 | 2.48 | 5.14 | 0.16 | 54.71 | 0.7 | 8.34 | 99.54 |
| SA3 | | 13.00 | 5.05 | 9.03 | 1.36 | 4.67 | 4.19 | 0.68 | 47.76 | 0.615 | 12.47 | 98.84 |
| SA4 | | 15.46 | 0.88 | 12.84 | 2.01 | 2.59 | 7.06 | 0.19 | 48.44 | 0.648 | 9.18 | 99.30 |
| SA5 | | 13.34 | 3.71 | 8.88 | 1.37 | 3.94 | 3.75 | 0.47 | 52.66 | 0.659 | 10.71 | 99.48 |
| SA6 | | 15.32 | 0.81 | 10.99 | 1.76 | 2.45 | 6.24 | 0.14 | 51.72 | 0.704 | 9.05 | 99.19 |
| SA7 | | 13.19 | 4.85 | 8.99 | 1.37 | 4.52 | 3.94 | 0.55 | 48.99 | 0.642 | 12.32 | 99.36 |
| SA8 | | 14.29 | 3.16 | 10.22 | 1.52 | 3.75 | 4.40 | 0.41 | 49.87 | 0.708 | 10.82 | 99.14 |
| **Mean** | | **14.22** | **3.00** | **10.07** | **1.55** | **3.58** | **4.79** | **0.42** | **50.39** | **0.67** | **10.60** | |
| **STD** | | **0.99** | **1.89** | **1.38** | **0.23** | **0.94** | **1.26** | **0.24** | **2.41** | **0.03** | **1.59** | |
| SB1 | SB group: Sediment from within Feature 1 | 14.85 | 0.82 | 8.99 | 1.58 | 1.52 | 4.53 | 0.14 | 58.77 | 0.652 | 7.41 | 99.25 |
| SB2 | | 14.49 | 1.20 | 9.92 | 1.42 | 2.18 | 4.31 | 0.46 | 56.52 | 0.682 | 7.8 | 98.99 |
| SB3 | | 16.65 | 0.72 | 8.83 | 1.50 | 1.68 | 3.48 | 0.09 | 57.65 | 0.713 | 7.97 | 99.28 |
| **Mean** | | **15.33** | **0.91** | **9.24** | **1.50** | **1.79** | **4.11** | **0.23** | **57.65** | **0.68** | **7.73** | |
| **STD** | | **1.16** | **0.25** | **0.59** | **0.08** | **0.35** | **0.55** | **0.20** | **1.13** | **0.03** | **0.29** | |

| | | | | | | | | | | | | |
|---|---|---|---|---|---|---|---|---|---|---|---|---|
| SC1 | SC group: Sediment between Features 1 and 2 | 15.32 | 0.76 | 8.25 | 1.69 | 1.38 | 2.68 | 0.14 | 61.98 | 0.767 | 6.8 | 99.77 |
| SC2 | | 17.10 | 0.54 | 9.30 | 1.74 | 1.59 | 3.24 | 0.10 | 56.77 | 0.827 | 8.24 | 99.45 |
| SC3 | | 12.03 | 0.69 | 7.45 | 1.42 | 1.37 | 3.72 | 0.10 | 65.79 | 0.561 | 6.18 | 99.30 |
| SC4 | | 15.43 | 0.95 | 10.57 | 1.64 | 1.86 | 5.24 | 0.35 | 53.63 | 0.717 | 9.16 | 99.55 |
| **Mean** | | **14.97** | **0.73** | **8.89** | **1.62** | **1.55** | **3.72** | **0.17** | **59.54** | **0.72** | **7.60** | |
| **STD** | | **2.12** | **0.17** | **1.35** | **0.14** | **0.23** | **1.10** | **0.12** | **5.40** | **0.11** | **1.35** | |
| SE1 | SE group: Sediment from vertical profile south of Feature 1 | 15.03 | 0.86 | 10.31 | 1.59 | 2.11 | 4.83 | 0.23 | 55.68 | 0.719 | 7.91 | 99.27 |
| SE2 | | 14.17 | 0.764 | 10.03 | 1.47 | 2.08 | 4.498 | 0.17 | 58.16 | 0.667 | 7.54 | 99.55 |
| SE3 | | 16.01 | 0.332 | 7.346 | 1.54 | 1.58 | 0.498 | 0.1 | 65.01 | 0.959 | 6.25 | 99.62 |
| SE4 | | 15.3 | 0.731 | 10.36 | 1.55 | 2.28 | 4.697 | 0.15 | 55.66 | 0.731 | 8.05 | 99.52 |
| SE5 | | 15 | 0.688 | 10.27 | 1.55 | 2.39 | 4.882 | 0.15 | 55.38 | 0.708 | 8.12 | 99.13 |
| **Mean** | | **15.10** | **0.68** | **9.66** | **1.54** | **2.09** | **3.88** | **0.16** | **57.98** | **0.76** | **7.57** | |
| **STD** | | **0.66** | **0.20** | **1.30** | **0.05** | **0.31** | **1.90** | **0.05** | **4.09** | **0.12** | **0.77** | |
| FS2280 | Puzzle Box excavation | 15.16 | 1.17 | 9.065 | 1.6 | 2.17 | 16.05 | 0.17 | 41.21 | 0.66 | 10.66 | 97.92 |

Supplementary Table 3: Trace element and rare earth elements (REEs) results. Separate Excel spreadsheet.

Supplementary Table 4: Instrumental parameters for WDXRFS (MagiX PRO). Measurements were carried out under vacuum with a rhodium X-ray tube without tube filter, a sample spinner, 25 mm collimator mask and a flowcounter. Elements were analysed in the order of decreasing X-ray energy; i.e., from Ni to Na.

| Element and X-ray line | Crystal | Collimator | Tube kV | Offset Bg1 (°2θ) | Offset Bg2 (°2θ) | PHD1 LL | PHD1 UL | Comment |
|---|---|---|---|---|---|---|---|---|
| Ni Kα | LiF 220 | 150 µm | 60 | -1.23 | 1.9 | 23 | 59 | |
| Fe Kα | LiF 220 | 150 µm | 60 | -3.25 | 5 | 18 | 60 | |
| Mn Kα | LiF 220 | 150 µm | 60 | -3.7 | 4.91 | 16 | 61 | |
| Cr Kα | LiF 220 | 150 µm | 50 | -2.4 | 2.95 | 14 | 60 | |
| V Kα | LiF 220 | 150 µm | 50 | -2.2 | 2.75 | 12 | 61 | |
| Ti Kα | LiF 200 | 150 µm | 40 | | | 36 | 62 | Used bg meas. of Ba. |
| Ba Lα | LiF 200 | 150 µm | 40 | -4.95 | 2.14 | 36 | 62 | |
| Ca Kα | LiF 200 | 150 µm | 30 | -4.7 | 3.55 | 36 | 62 | |
| K Kα | LiF 200 | 150 µm | 30 | -5.3 | 5.5 | 36 | 63 | |
| S Kα | Ge 111 | 550 µm | 30 | 5.7 | | 35 | 64 | Bg factor 1.16. |
| P Kα | Ge 111 | 550 µm | 30 | -10.24 | 6.5 | 35 | 64 | |
| Si Kα | PE 002 | 550 µm | 30 | -4.2 | 5.69 | 33 | 67 | |
| Br $L\beta_1$ | PE 002 | 550 µm | 30 | | | 33 | 67 | Used bg of Al. |
| Al Kα | PE 002 | 550 µm | 30 | -11.28 | | 32 | 67 | Bg factor 1.0. |
| Mg Kα | PX1 | 150 µm | 30 | -2 | | 34 | 66 | Used bg1 meas. of Na. |
| Na Kα | PX1 | 150 µm | 30 | -2.384 | 2 | 32 | 68 | |

Notes: LL and UL are the lower and upper level, respectively, of the pulse height analyser window. PX1 is a synthetic multilayer with a nominal 2d spacing of 5 nm.

Supplementary Table 5: Instrument parameters for ICPMS (NexION, Perkin-Elmer)

| | |
|---|---|
| Sample introduction system | Cross-flow with Scott double-pass spray chamber |
| Nebuliser gas flow | Ca. 0.8 to 0.9 L/min |

| | |
|---|---|
| Sample uptake rate | Ca. 1 mL/min |
| Skimmer cones | Nickel |
| RF power | 1200 W |
| Data acquisition | Peak hoping mode, 20 sweeps per reading, 1 reading per replicate, 3 replicates. Dwell time 40 ms. |

### Supplementary Information: Figures

**Supplementary Figure S1.** Geological face map and cross-sections through the sediments at different locations in the Dinaledi Chamber, illustrating the relationships between the flowstone groups and sedimentary units. Figure modified from Dirks and colleagues (2017: figure 2) to remove hypothesized subsurface floor drain, discussed in text.

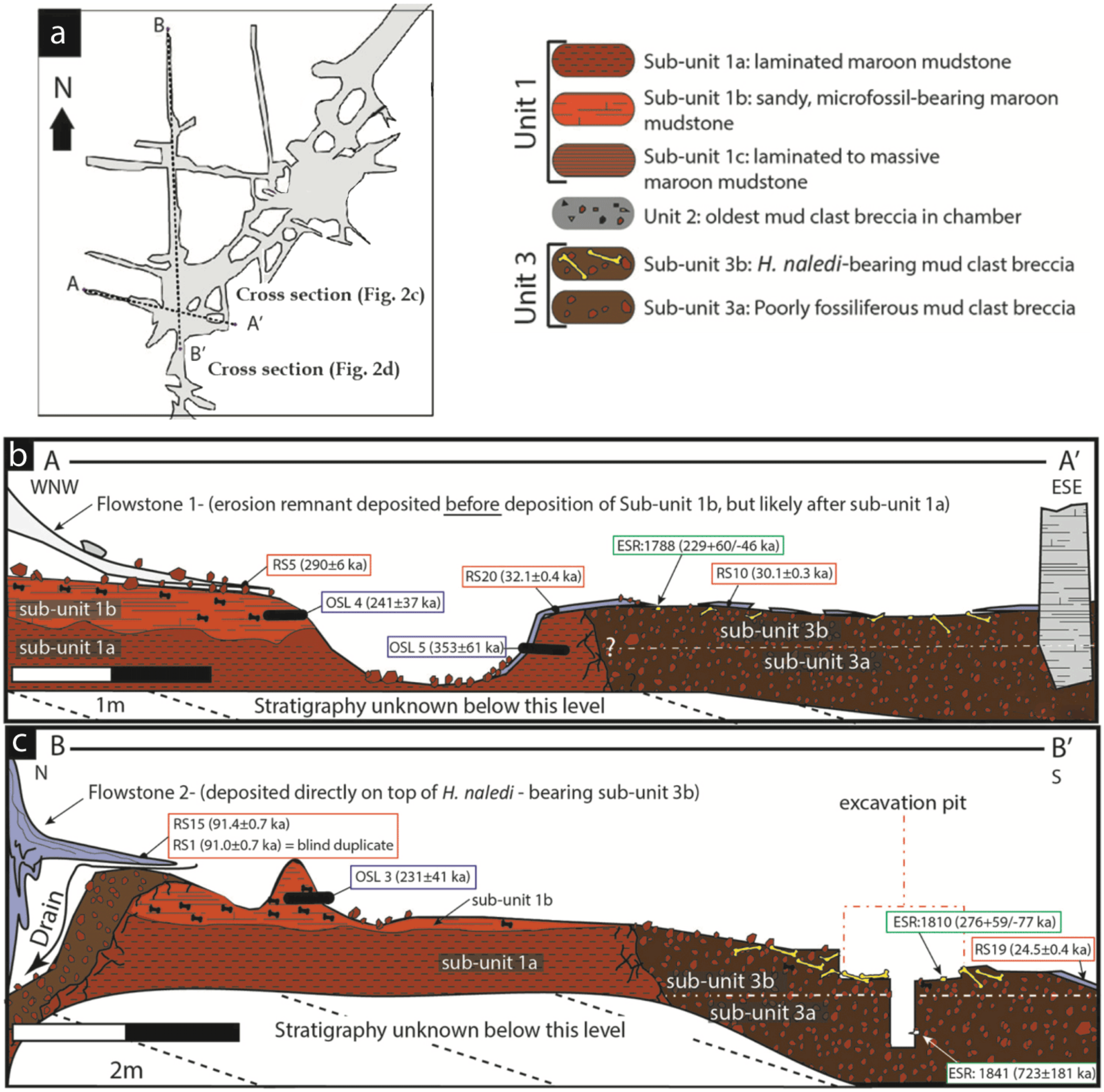

**Supplementary Figure S2.** Data and characteristics of cave floor sediments (Facies 2) from the Dinaledi and Dragon's Back Chambers. Figure taken from Dirks and colleagues (2015: figure 5 - https://doi.org/10.7554/eLife.09561.007). Original figure legend: (A) Grain size distribution of sample UW101-SO-39 (Figure 2C). The bulk of the sample material falls within a size fraction corresponding to silt and fine-grained sand. Some coarser mudstone fragments did not disintegrate when immersed in water, likely due to considerable Mn- and Fe-oxide micro-concretionary development in the orange mudstone. Because some mudstone fragments are well lithified the particle size distribution is skewed towards the coarser grain-size values. (B) Results of XRF analyses of bulk samples of three floor sediments from the Dinaledi Chamber (UW101-SO31, -34 and -39) and one from the Dragon's Back chamber (DB-1). The sample from the Dragon's Back Chamber has a radically different composition from those of the Dinaledi Chamber, with the high $SiO_2$ content reflecting its dominance of quartz. The Dinaledi samples have much higher $Al_2O_3$ and $K_2O$ contents than DB-1, indicating a higher content of clay minerals and mica, and higher CaO, MgO, MnO, and total Fe oxide contents which reflect alterations and inclusions. The higher $P_2O_5$ content of the Dinaledi samples is probably located in comminuted bone fragments which are seen macroscopically. The volatiles content (LOI) of the Dinaledi samples is also higher than in DB-1, in accord with a higher total clay mineral and mica content. (C–E) Backscattered electron (BSE) wide-field images of grain mounts from floor sediments. Brighter shades indicate the presence of heavier elements, mainly Mn and Fe in altered grains. (C) DB-1, Dragon's Back Chamber, large fragments are quartz and chert, partly altered. (D) UW101-SO34. (E) UW101-SO39. In these samples the large fragments are almost exclusively clay; note their angular shape which shows these to be locally derived.

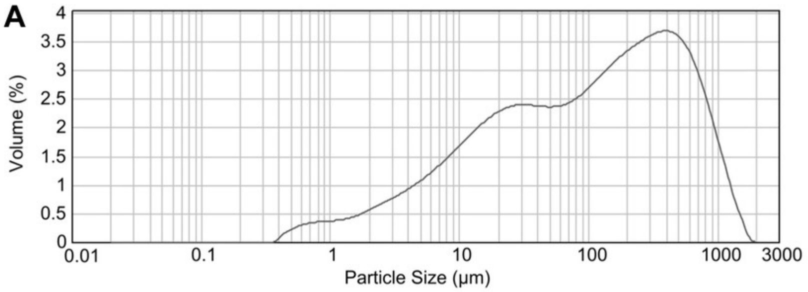

B

| | DB1 | UW 101-SO31 | UW 101-SO34 | UW 101-SO39 |
|---|---|---|---|---|
| $SiO_2$ | 84.68 | 51.33 | 51.16 | 51.79 |
| $TiO_2$ | 0.37 | 0.7 | 0.62 | 0.7 |
| $Al_2O_3$ | 4.33 | 15.85 | 13.96 | 16.2 |
| MnO | 0.88 | 3.92 | 4.26 | 4.32 |
| $Fe_2O_3$ | 3.73 | 10.43 | 10.7 | 10.95 |
| MgO | 0.46 | 3.21 | 4.01 | 3.1 |
| CaO | 0.62 | 2.13 | 2.4 | 1.06 |
| $K_2O$ | 0.48 | 1.52 | 1.35 | 1.61 |
| $Na_2O$ | 0.06 | 0.1 | 0.09 | 0.08 |
| $P_2O_5$ | 0.16 | 0.97 | 0.48 | 0.29 |
| BaO | - | 0.06 | 0.06 | 0.06 |
| $SO_3$ | - | 0.09 | 0.1 | - |
| LOI | 3.27 | 9.02 | 9.84 | 9.04 |
| Sum | 99.04 | 99.33 | 99.03 | 99.2 |

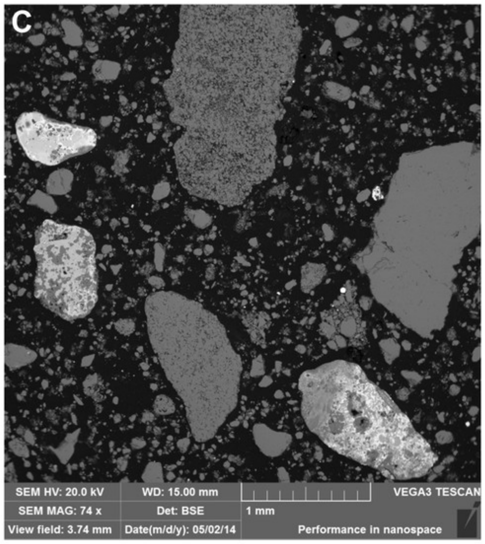

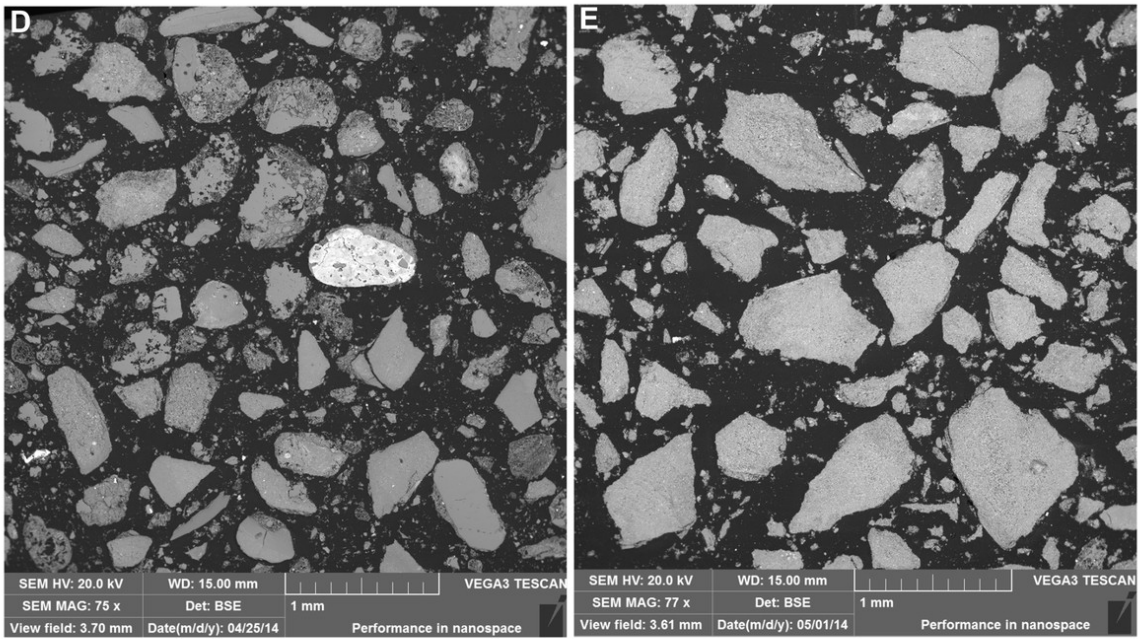

**Supplementary Figure S3.** Stratigraphic units and flowstones observed in the Dinaledi Chamber, showing Unit 1 sediment lamination and laminated orange-red mud clasts. Figure taken from Dirks and colleagues (2015: figure 4 - https://doi.org/10.7554/eLife.09561.006). Original figure legend: (A) Erosional remnant of horizontally laminated Unit 1 strata (Facies 1). (B) Close-up view of Unit 1 (Facies 1a) showing fine laminations and small invertebrate burrows (note fine sand infilling in burrows). (C) Overview photo of the Dinaledi Chamber, directly to the east of the entrance point into the chamber. Photo shows distribution of Flowstones 1–3 and stratigraphic Units 2 and 3. (D) Close-up view of Flowstone 1 encasing sediment of Unit 2. Note that several generations of flowstone (Flowstones 1a–e) are coating Unit 2. The thin, clear lower layer is Flowstone 1a, and the overlying white flowstone is either Flowstone 2 or 3. (E) Close-up view of Unit 2, consisting of generally poorly-cemented Facies 2 sediment. (F) View of the chamber floor near the entry point. On the cave floor, a large erosional remnant of Unit 1 (orange laminated mudstone of Facies 1a), is surrounded by mud-clast breccia of Unit 3 (main hominin bearing unit). Note that Flowstone 2 has been undercut by post-depositional erosion of Unit 3, which, in this location has resulted in a lowering of the floor by as much as 25 cm. (G) Flowstone 2 overlying Unit 3 in one of the chamber's side passages. In this location Unit 3 has also been partly eroded after depositional from underneath the flowstone drape, leaving a hanging remnant, with some indurated sediment of Unit 3 attached to its base. Note the continued deposition of sediment above Flowstone 2.

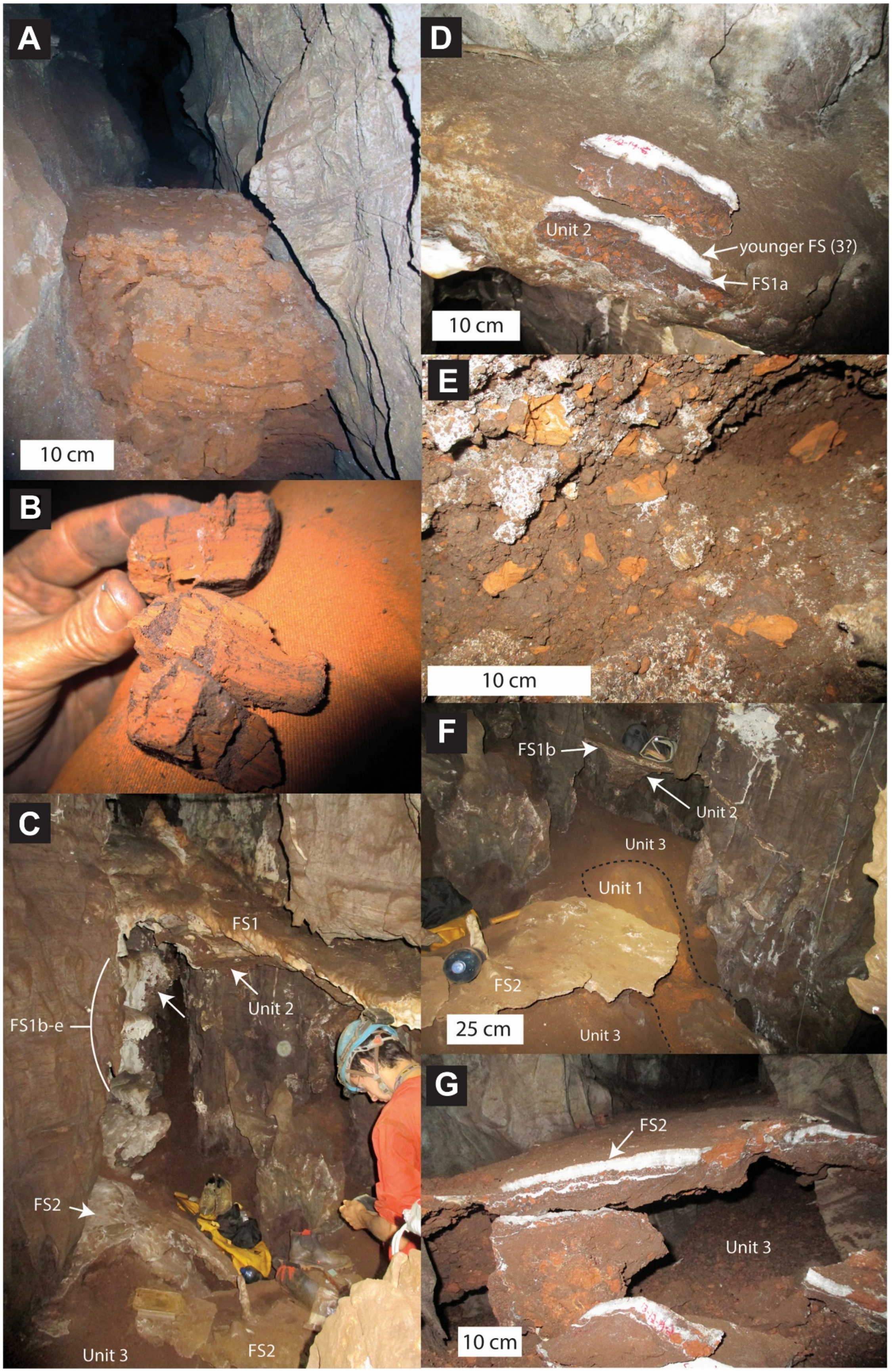
A
10 cm
B
C
FS1
FS1b-e
Unit 2
FS2
Unit 3
FS2
D
Unit 2
younger FS (3?)
FS1a
10 cm
E
10 cm
F
FS1b
Unit 2
Unit 3
Unit 1
FS2
25 cm
Unit 3
G
FS2
Unit 3
10 cm

**Supplementary Figure S4.** Hill Antechamber excavation unit S150W150 prior to opening excavation. This area had a collection of non-overlapping flat stones on the surface.

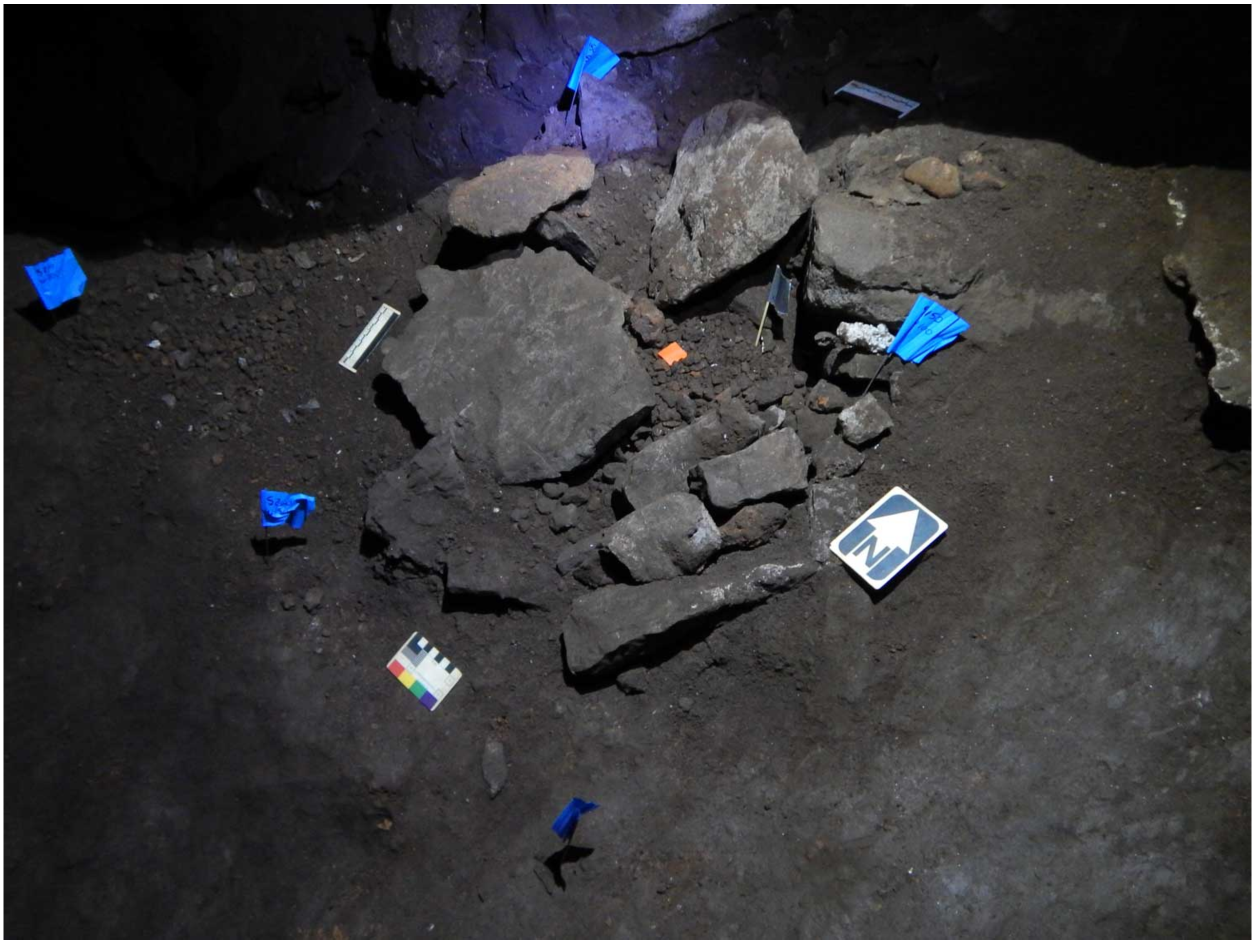

**Supplementary Figure S5.** Top surface of the Hill Antechamber feature after full exposure. Photo (left) and 3D model based on photogrammetry (right). The bone material at the northmost extent of the feature is powdery and highly fragmented, with more complete skeletal elements visible toward the south.

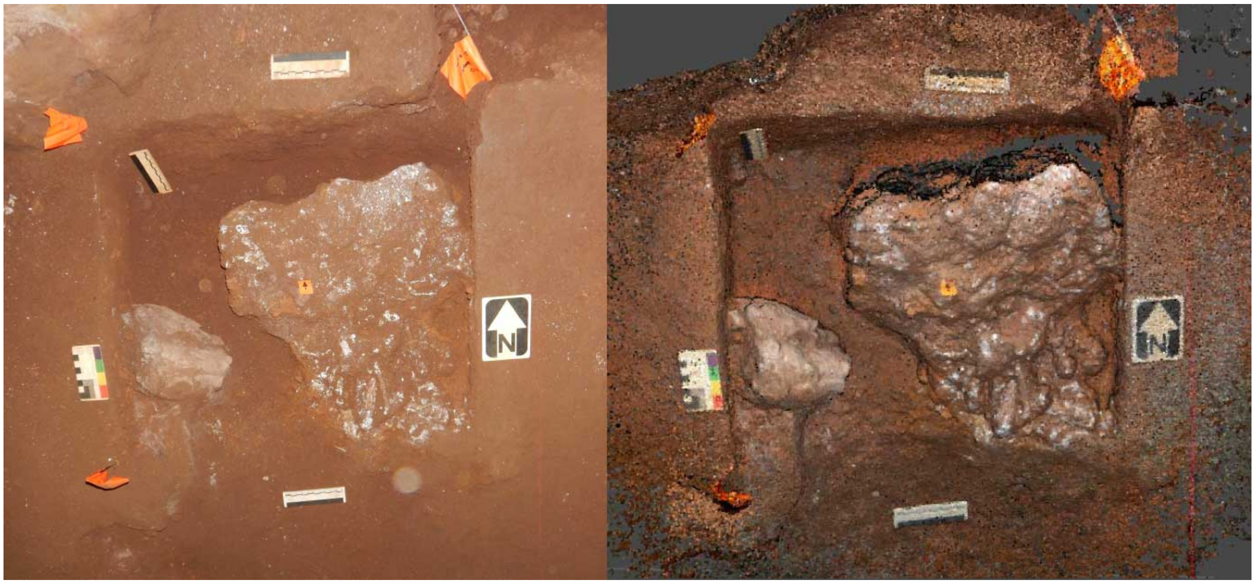

**Supplementary Figure S6.** Hill Antechamber feature after pedestaling and separation from surrounding sediment.

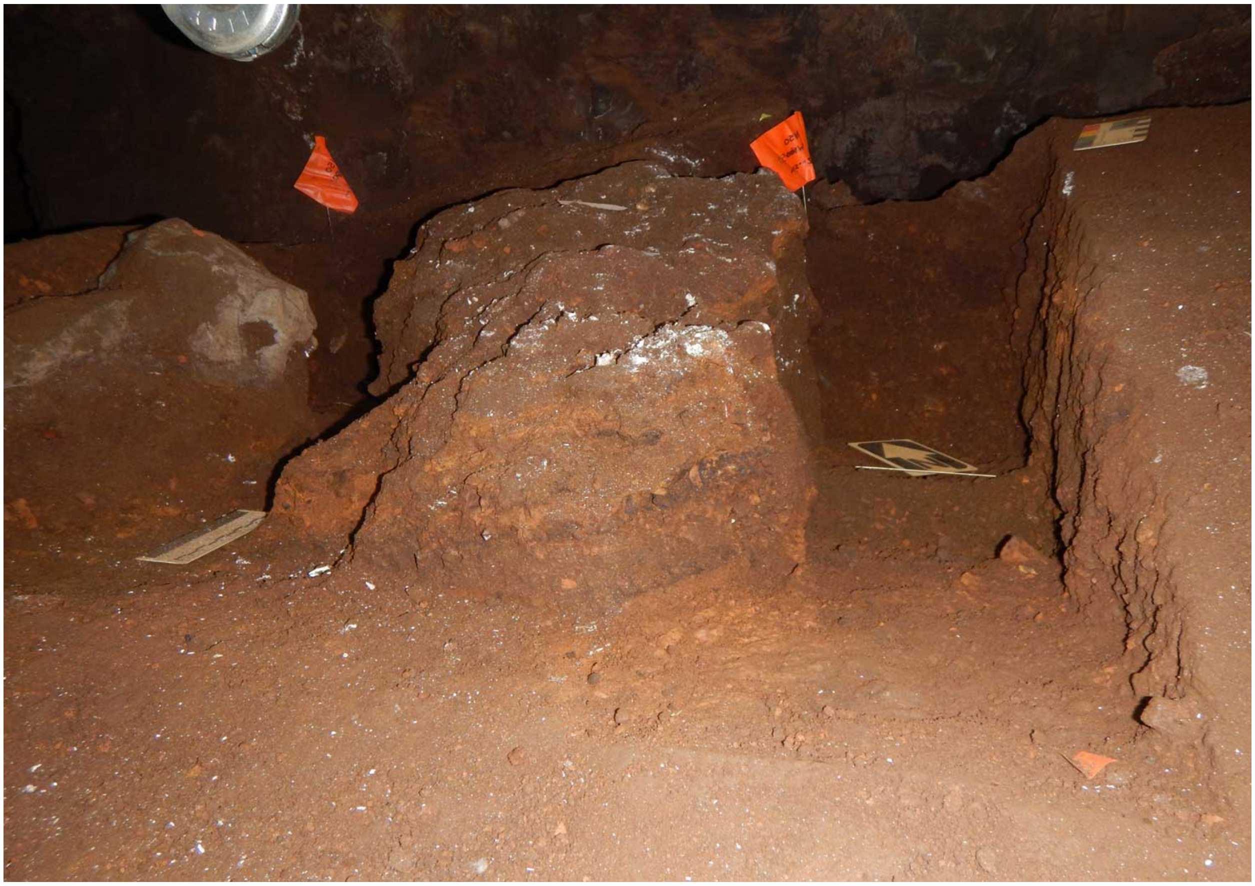

**Supplementary Figure S7.** Sagittal (north-south) section of Hill Antechamber feature. North is at left of frame. This section is at approximately 55% of the east-west breadth of the feature. The articulated foot is visible in longitudinal section at right of frame, with cross-sections of other bones and teeth further to the left of frame. The layer that constitutes the top of the feature is packed with bone material including articulated, semi-articulated and loose material, flattened into less than 5 cm thickness.

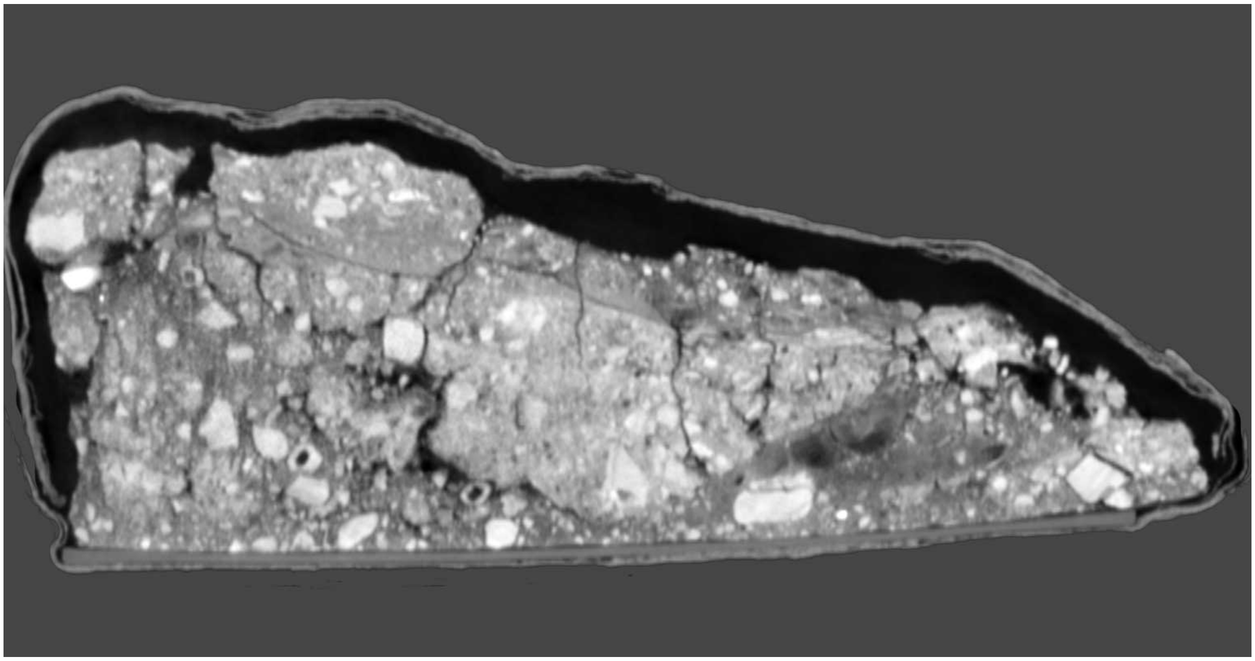

**Supplementary Figure S8.** Hill Antechamber feature after jacketing of largest block (U.W. 101-2076) in six layers of plaster bandages, prior to separation from sediment at its base.

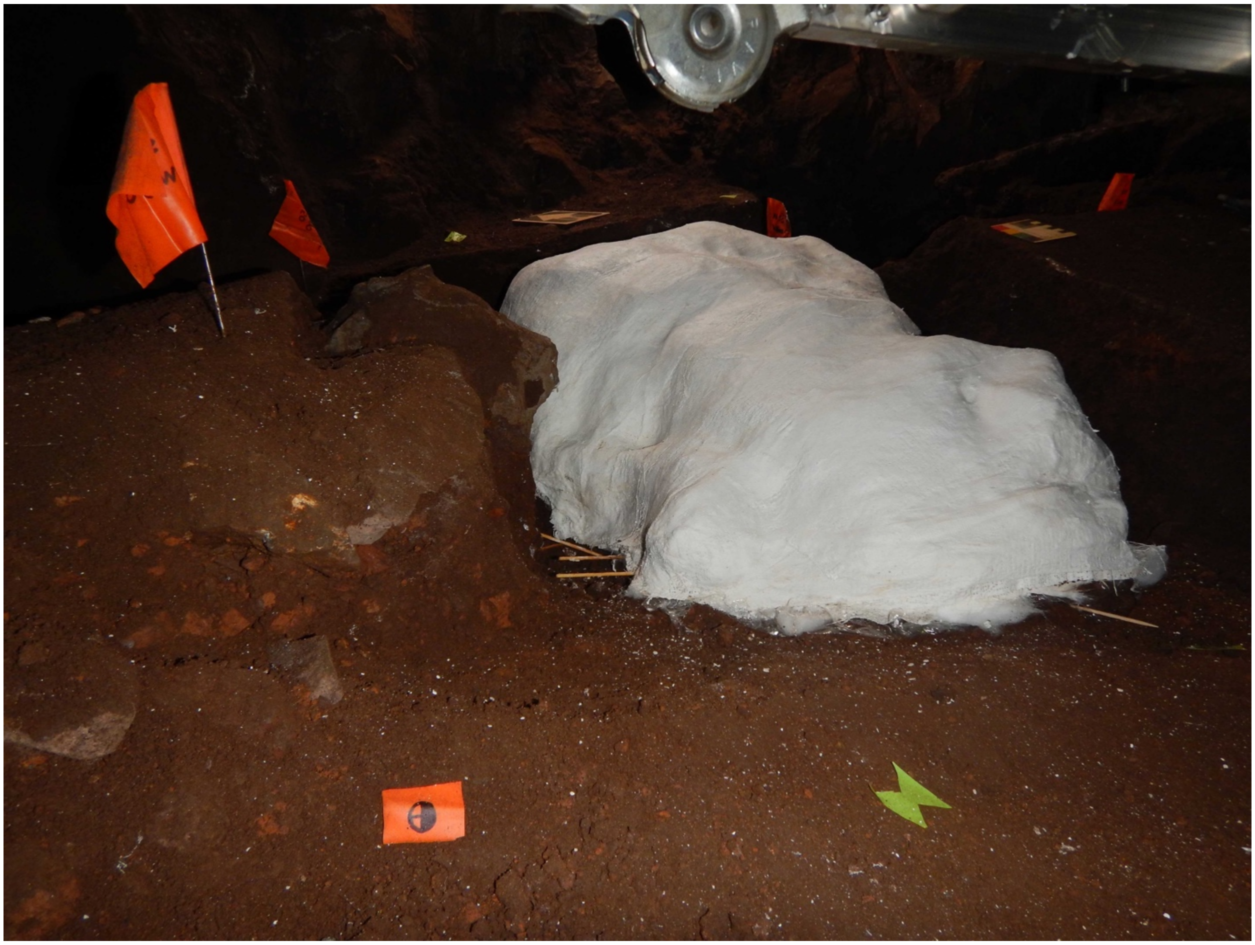

**Supplementary Figure S9.** Fossil mass within plaster jacket after packing into waterproof caving bag for exit from the cave system.

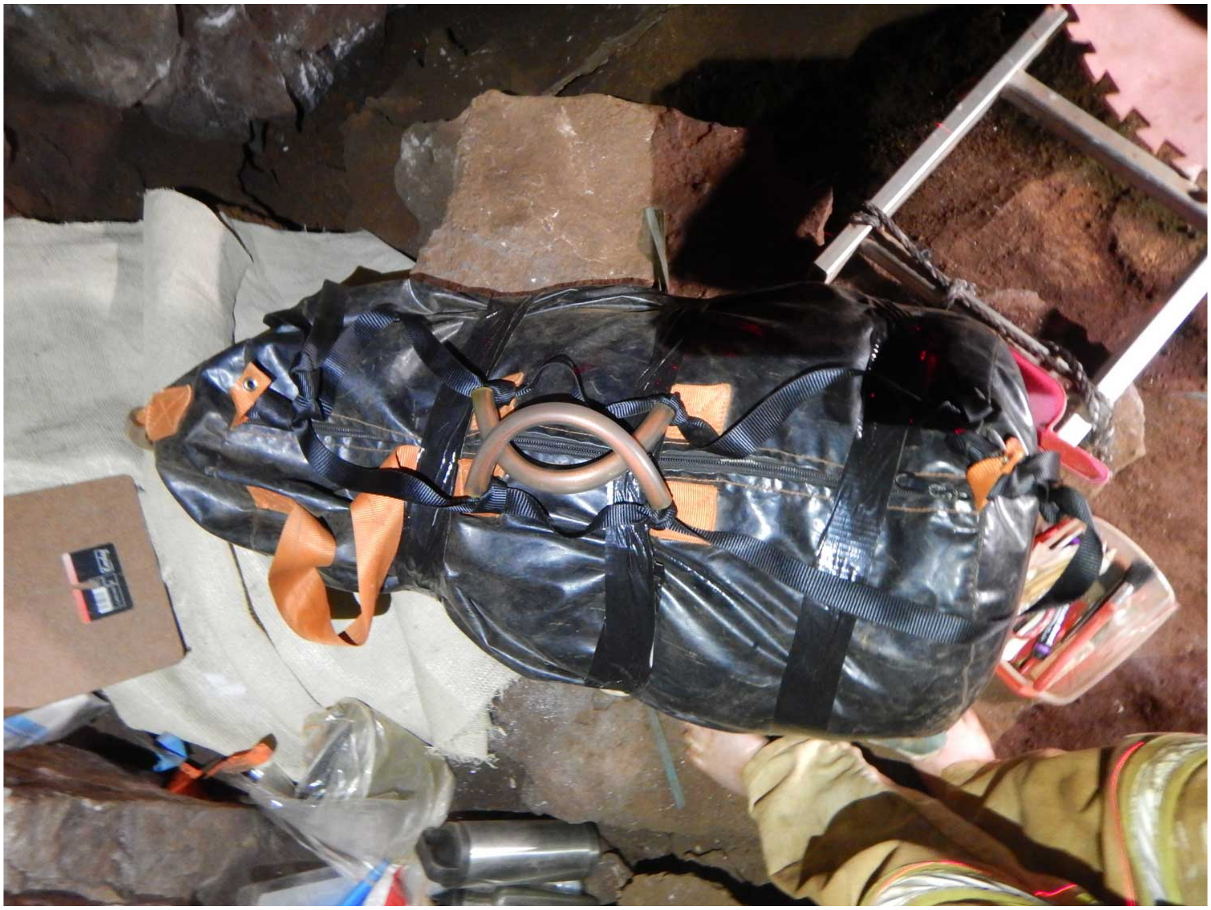

**Supplementary Figure S10.** Detail of medical-resolution CT image of Hill Antechamber feature. This is a horizontal section with north at bottom of frame and west at right of frame. In this image, the bright object at lower left is a cross section of HAA1. At its left, cross sections of four rays of the articulated hand are visible; there is also a bone visible in the gap or space adjacent to the artifact that is a fragment of intermediate phalanx.

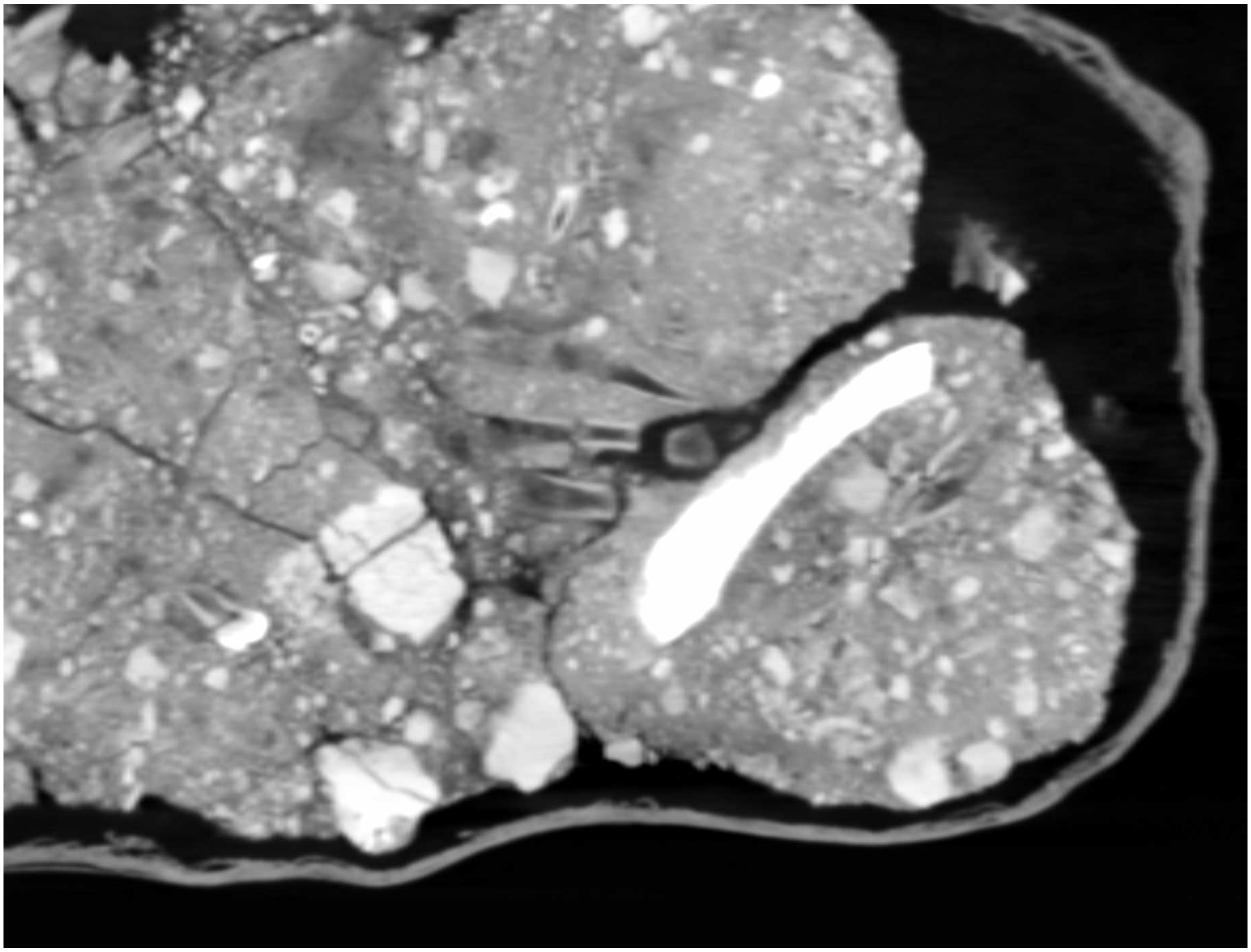

**Supplementary Figure S11.** Sediment profile showing east wall of N100W50 excavation unit in Hill Antechamber. The sediment is a dark brown unlithified breccia containing laminated orange-red mud (LORM) clasts. In this unit, the clasts make up a small fraction of the sedimentary deposit with little evidence of layering or stratigraphic differentiation.

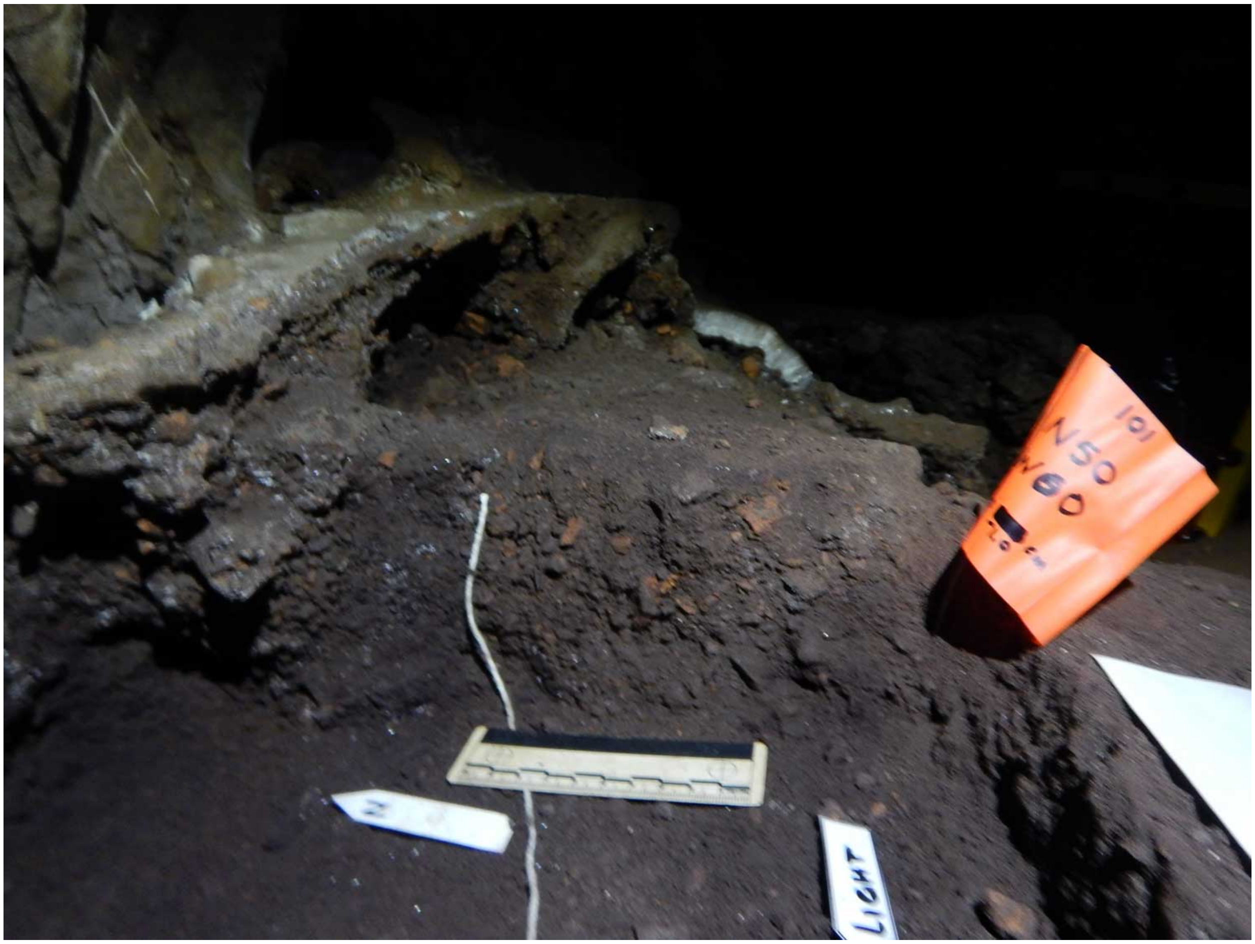

**Supplementary Figure S12.** Hill Antechamber excavation east wall profile. North is at left of frame. The ellipse is drawn around a 15 cm by 10 cm by 5 cm collapse of the wall that accompanied removal of the plaster jacketed block, resulting in some distortion to the profile in this localized area. The darker patch at the right side of the ellipse is a shadow from the collapse edge, not a dark-colored inclusion in the sediment. Layering of the unlithified mud clast breccia is visible with some layers having a higher content of LORM clasts and laminae, with color variation less evident here than in the section shown in Figure S11 and S13 The layering is approximately parallel to the slope of the chamber floor. LORM content and clasts are less toward the north edge of the excavation, at left of frame.

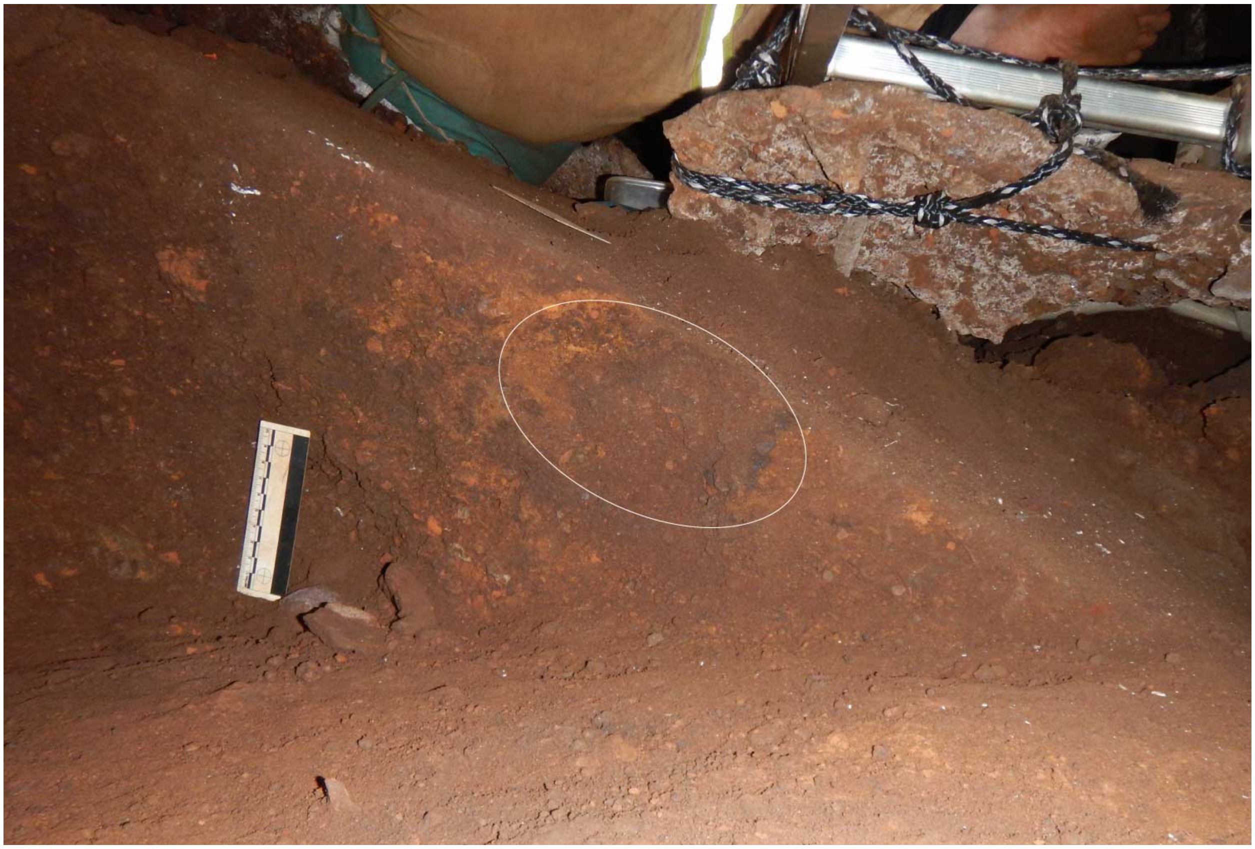

**Supplementary Figure S13.** Stratigraphic profile of sediments directly adjacent to south boundary of Hill Antechamber feature. At left of image is the west side of the unit. This profile represents the sedimentary structure of the S50W100 and S50W50 units before the excavation reduced the feature to its rounded south edge. Horizontal layering of unlithified mud clast breccia (UMCB) with denser orange-red LORM-clast bearing laminae is evident for the top 10 cm of the profile. This layering becomes subhorizontal with horizontal depth with an east-west trending slope. The lowest layer visible trends into the horizontal floor of the excavation unit. This layering is not paralleled by the skeletal material, fill, or LORM clasts visible within the feature itself.

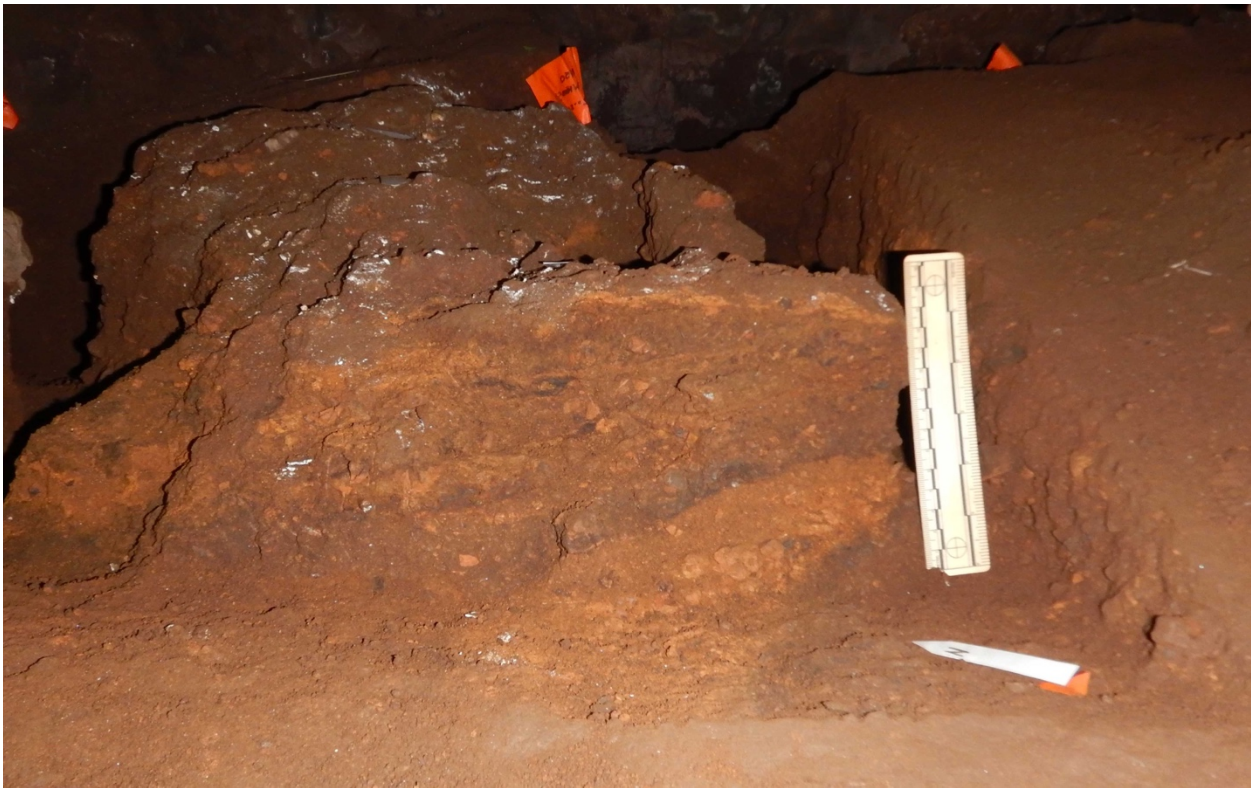

**Supplementary Figure S14.** CT section of Hill Antechamber feature. This east-west transverse section is at approximately 50% of north-south length of the feature. At the bottom of the section, many small LORM clasts are visible, with two notable voids taking the form of vertical cracks. The disordered array of LORM clasts continues to the right of image with frequent voids (west side of feature).

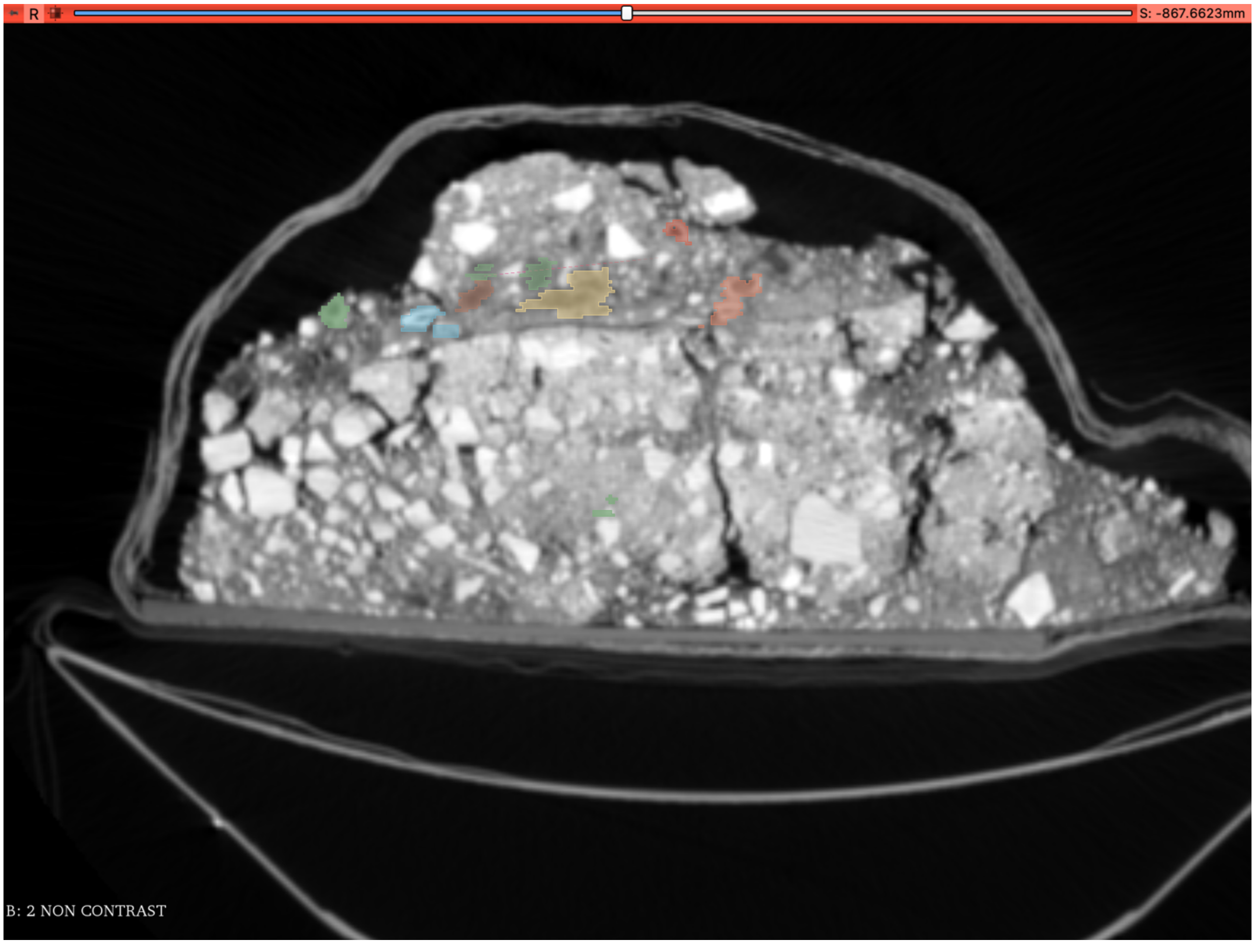

**Supplementary Figure S15.** CT section of Hill Antechamber feature. This is a transverse section on the east-west plane at approximately 65% of the north-south length. The five rays of the articulated foot are visible at lower left of the section. This section cuts across the metatarsals. The bones of the foot are immediately surrounded by a halo of sediment that approximates the shape of the foot's soft tissue. This lower-density sediment separates the bones of the foot from surrounding, more radio-opaque LORM clasts and sediment. Above the foot some small voids in the sediment are visible; small voids are also visible toward the left of this section directly above a disordered arrangement of LORM clasts.

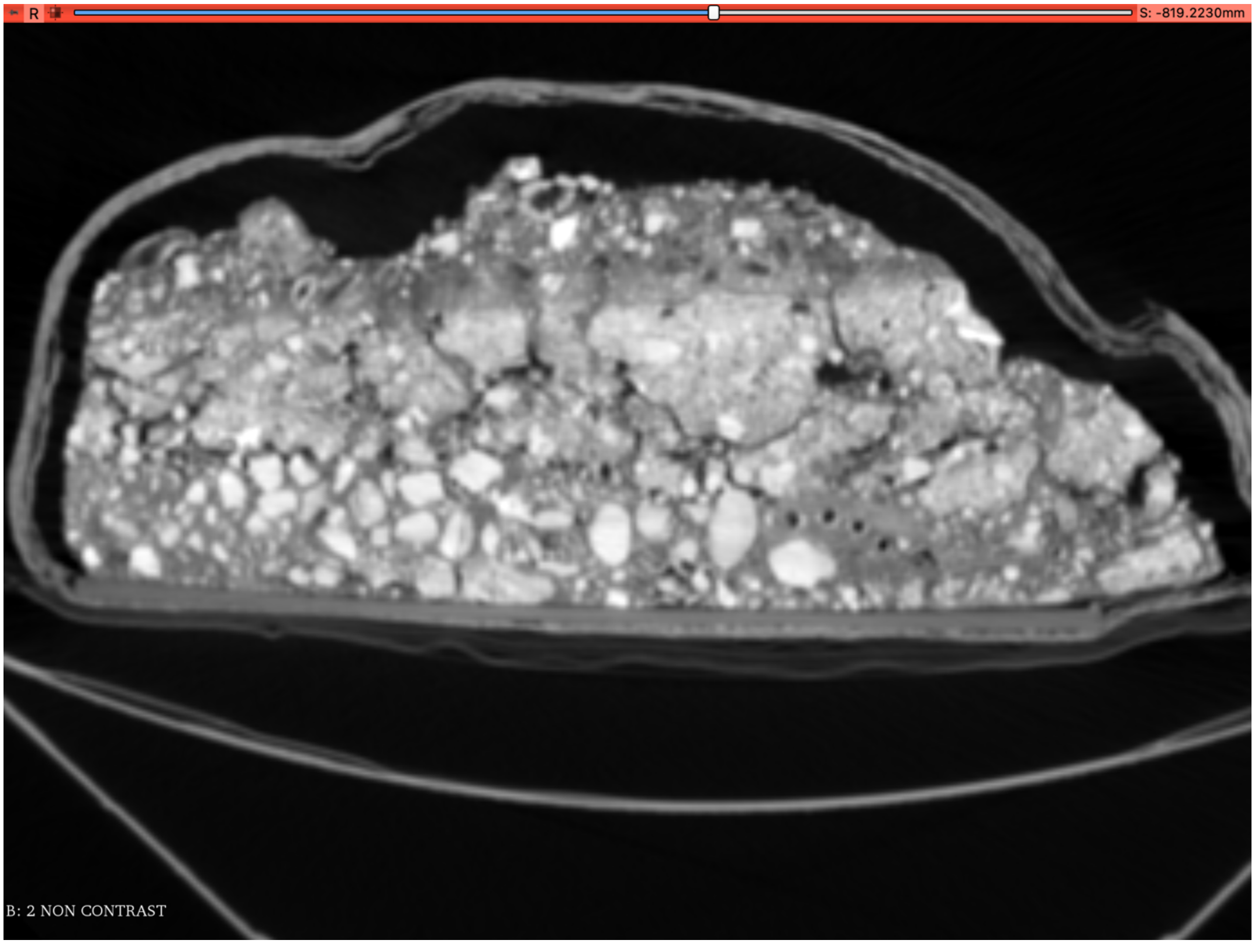

**Supplementary Figure S16.** Bright orange LORM patch immediately beneath Hill Antechamber feature. (A) Bottom of Hill Antechamber feature after jacketed extraction and inversion. The bright orange patch is visible centrally slightly toward the right (west) side of the inverted feature (B) Excavation unit immediately after jacketed extraction of feature and cleaning of surface. The corresponding orange patch is visible at center of unit.

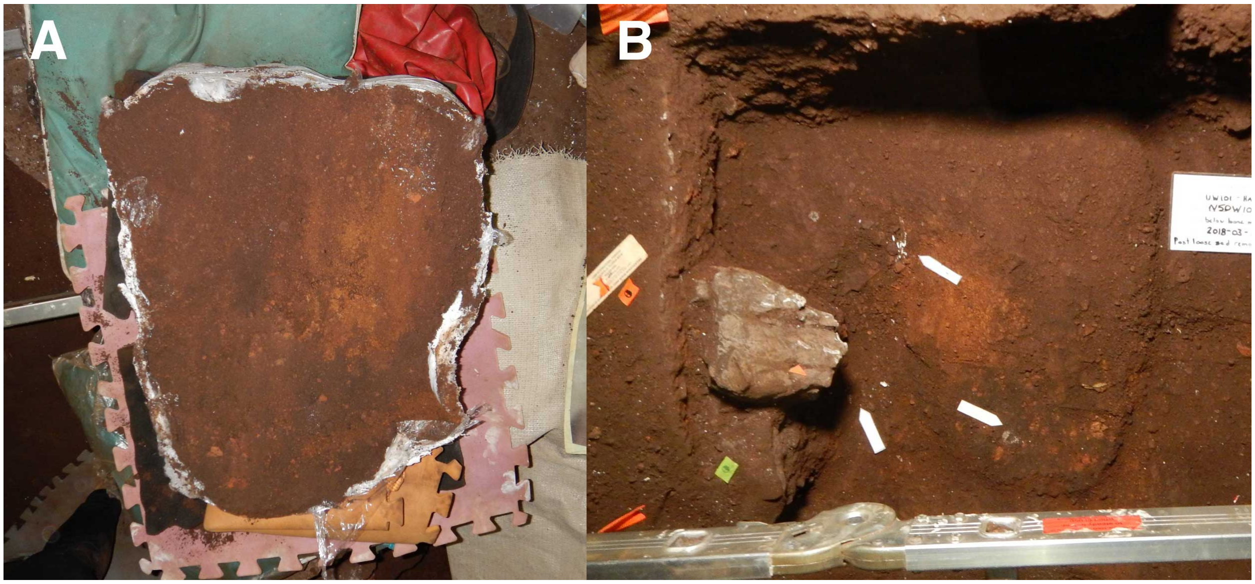

**Supplementary Figure S17.** Grain size distribution curves of the five sediment groups analyzed within and around Feature 1.

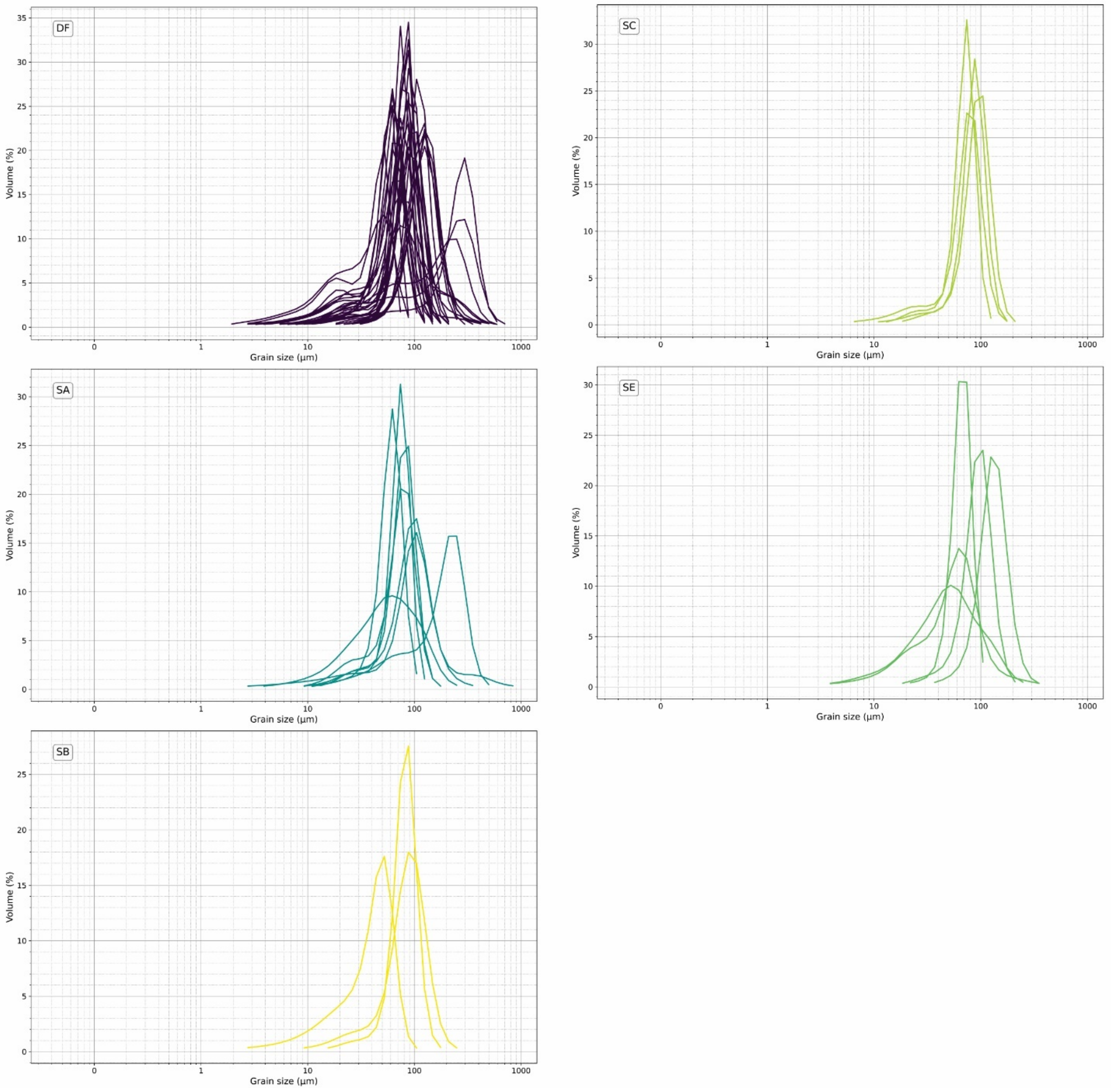

**Supplementary Figure S18.** Harker variation plots illustrating the relationship between CaO and selected rare earth elements (REEs) for the five sediment groups analyzed within and around Feature 1.

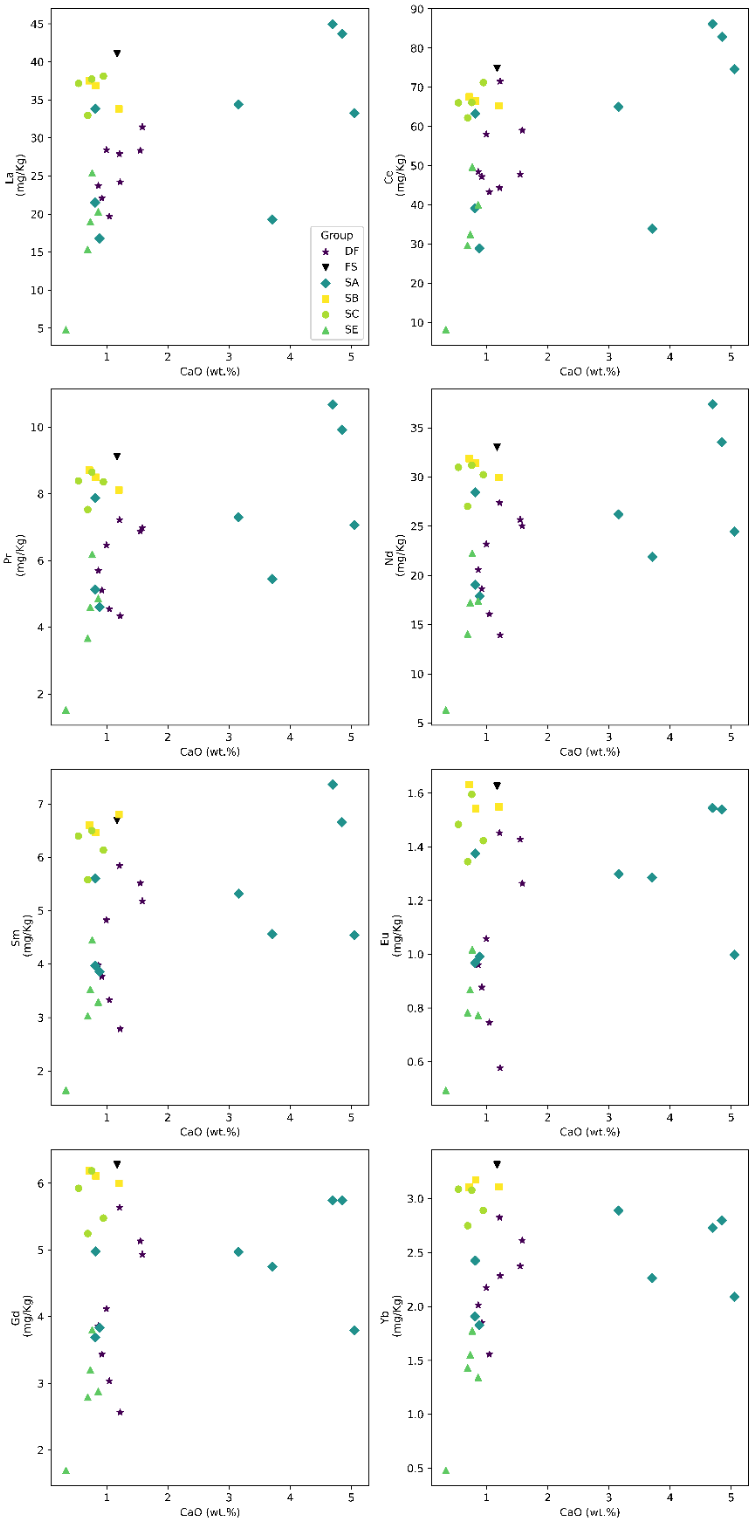

**Supplementary Figure S19.** Harker variation plots illustrating the relationship between Zn and selected rare earth elements (REEs) for the five sediment groups analyzed within and around Feature 1. SB plots distinctly apart from other sediment groups, which tend to overlap or are mingled for most trace elements.

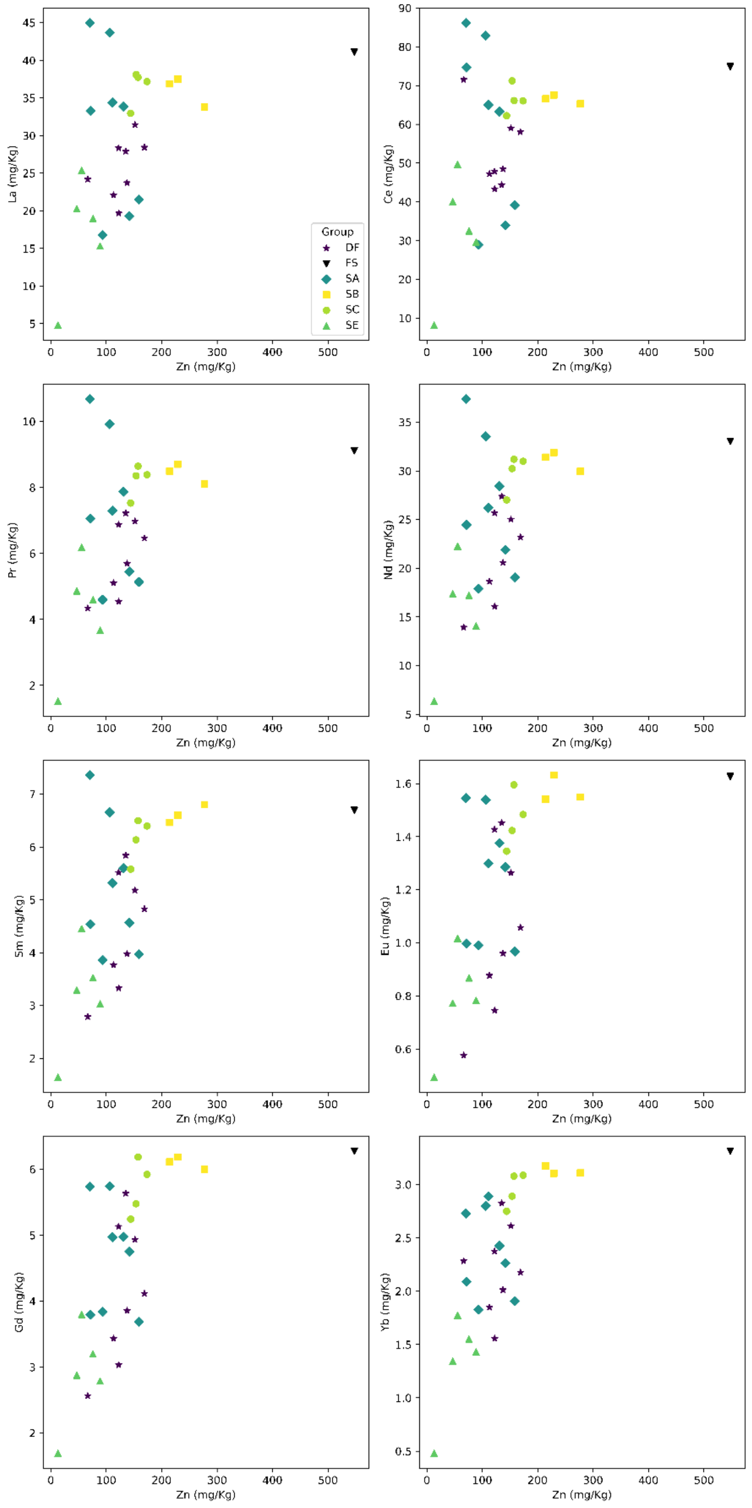

**Supplementary Figure S20.** Reconstruction of the burial position of Dinaledi Hand 1 and Foot 1. Hand 1 (blue) is nearly complete and shows the flexed (curled) nature of the fingers upon recovery. Foot 1 (purple) is less well preserved, but demonstrates the retention of the structures of the mid and hind foot. F1 is underlain by disarticulated manual elements not associated with Hand 1.

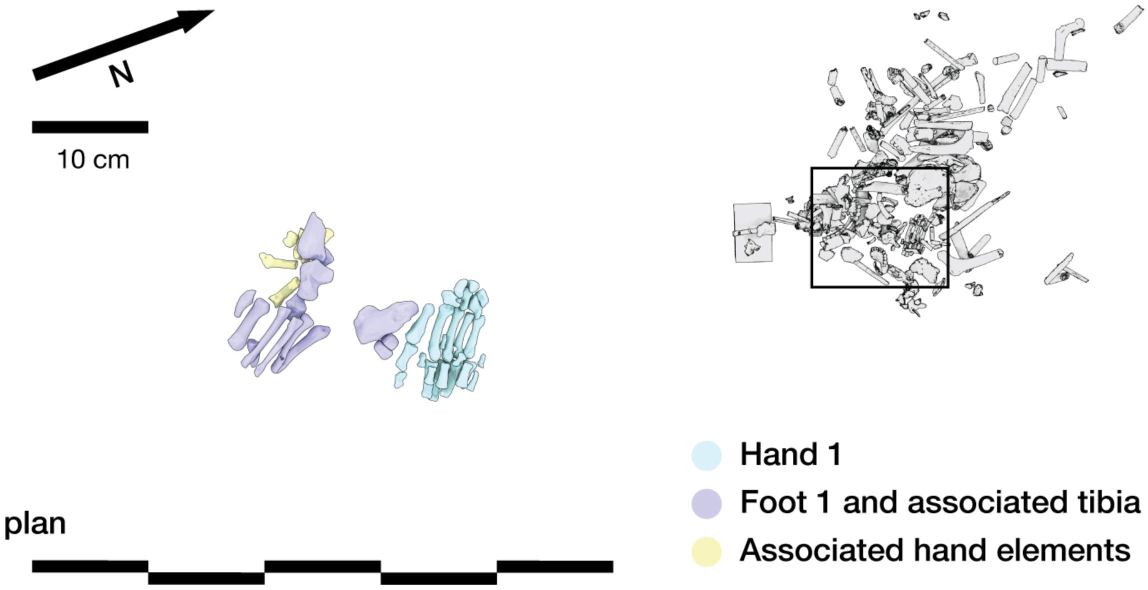

**Supplementary Figure S21.** Hill Antechamber Artifact 1 (HAA1) showing surface from 8 different angles with 2 different lighting directions. The 3D model results from the segmentation of the synchrotron scan at 16.22um, the artifact being still in situ in the paster jacket.

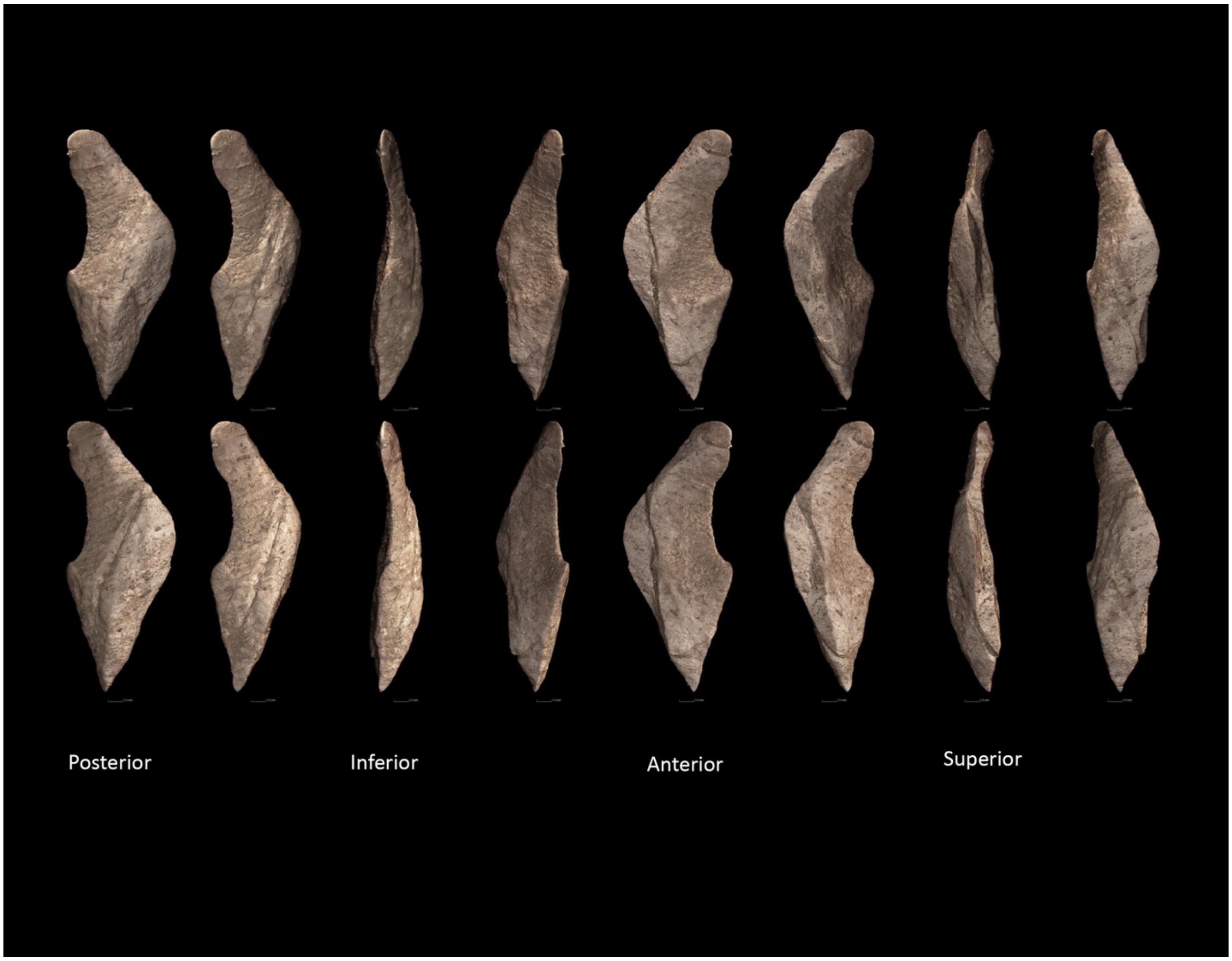

**Supplementary Figure S22.** Hill Antechamber Artifact 1 (HAA1) close-up from the previous figure with detail showing striations visible on both faces and intersection of these striations with sharp edge of artifact showing appearance of serrations.

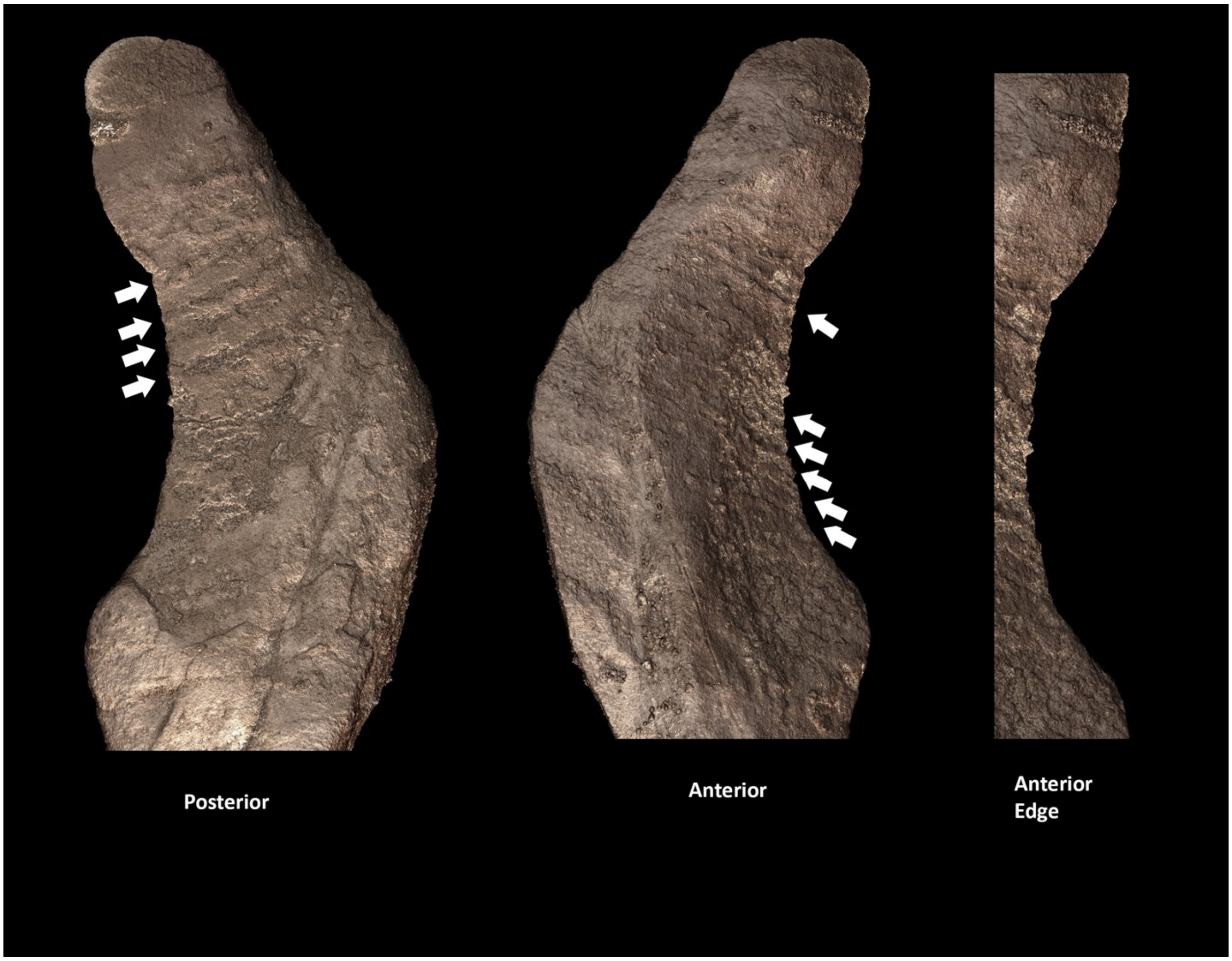
